# ZNF410 represses fetal globin by devoted control of CHD4/NuRD

**DOI:** 10.1101/2020.08.31.272856

**Authors:** Divya S. Vinjamur, Qiuming Yao, Mitchel A. Cole, Connor McGuckin, Chunyan Ren, Jing Zeng, Mir Hossain, Kevin Luk, Scot A. Wolfe, Luca Pinello, Daniel E. Bauer

## Abstract

Major effectors of adult-stage fetal globin silencing include the transcription factors (TFs) BCL11A and ZBTB7A/LRF and the NuRD chromatin complex, although each has potential on-target liabilities for rational *β*-hemoglobinopathy therapeutic inhibition. Here through CRISPR screening we discover ZNF410 to be a novel fetal hemoglobin (HbF) repressing TF. ZNF410 does not bind directly to the *γ*-globin genes but rather its chromatin occupancy is solely concentrated at *CHD4*, encoding the NuRD nucleosome remodeler, itself required for HbF repression. *CHD4* has two ZNF410-bound regulatory elements with 27 combined ZNF410 binding motifs constituting unparalleled genomic clusters. These elements completely account for ZNF410’s effects on *γ*-globin repression. Knockout of ZNF410 reduces CHD4 by 60%, enough to substantially de-repress HbF while avoiding the cellular toxicity of complete CHD4 loss. Mice with constitutive deficiency of the homolog Zfp410 are born at expected Mendelian ratios with unremarkable hematology. ZNF410 is dispensable for human hematopoietic engraftment potential and erythroid maturation unlike known HbF repressors. These studies identify a new rational target for HbF induction for the *β*-hemoglobin disorders with a wide therapeutic index. More broadly, ZNF410 represents a special class of gene regulator, a conserved transcription factor with singular devotion to regulation of a chromatin subcomplex.

## Introduction

Despite renewed enthusiasm for novel approaches to *β*-hemoglobinopathies, the clinical unmet need for these most common monogenic diseases remains vast^1–3^. Induction of fetal *γ*-globin gene expression could bypass the underlying *β*-globin molecular defects and ameliorate the pathophysiological cascades that result in elevated morbidity and mortality. Critical regulators of the switch from fetal to adult globin gene expression include the DNA-binding transcription factors (TFs) BCL11A and ZBTB7A and the nucleosome remodeling and deacetylase (NuRD) chromatin complex^4–7^. BCL11A and ZBTB7A each bind to unique sites at the proximal promoters of the duplicated fetal *γ*-globin genes *HBG1* and *HBG2* and each physically interact with NuRD^5,8–10^. Although the molecular details underpinning this switch, including the precise sequences bound and NuRD subcomplex members required, are increasingly understood, still the feasibility to directly perturb these mechanisms through pharmacology remains uncertain. One challenge is the pleiotropic molecular, cellular and organismal effects of each of the aforementioned fetal hemoglobin (HbF) repressors which makes the therapeutic window uncertain and the risk of undesired on-target liabilities considerable. An ideal target would have a wide therapeutic index through which inhibition of function could be tolerated across a diverse set of cellular contexts.

To better define additional molecular players orchestrating the developmental regulation of globin gene expression, we performed a CRISPR screen focusing on putative DNA-binding TFs that contribute to HbF silencing. We identified ZNF410 as a novel DNA-binding TF required for HbF repression. Little was previously known about ZNF410. We show indeed this gene is required for HbF silencing. Surprisingly we find that it displays a narrowly restricted pattern of chromatin occupancy, not binding to the globin locus directly, but rather binding to upstream elements, through an unusual set of clustered motifs, controlling the expression of the catalytic NuRD subunit CHD4. We observe that ZNF410 and its mouse homolog Zfp410 are dispensable for survival to adulthood as well as normal erythropoiesis and hematopoietic repopulation.

## Results

### CRISPR screen for novel transcriptional regulators of HbF level

We performed a CRISPR screen in a primary human erythroid precursor cell line (HUDEP-2) that expresses an adult-type pattern of globins to discover genes required for repression of HbF. The screen targeted 1591 transcription factors and 13 genes of the NuRD complex as controls. HUDEP-2 cells stably expressing SpCas9 were first generated. HUDEP-2/Cas9 cells were then transduced by the sgRNA library at low multiplicity and selected for sgRNA cassette integration by acquisition of puromycin resistance. Following erythroid maturation culture, cells were stained for HbF expression and HbF+ cells (range 1.8-7%) selected by FACS (**Fig. 1a**). Integrated sgRNAs were amplified from genomic DNA and counted by deep sequencing. We calculated two sgRNA enrichment scores. First, sgRNA abundance was compared in HbF+ and total cells at the end of erythroid maturation to obtain an HbF enrichment score. Second, sgRNA abundance was compared in cells at the end of erythroid maturation and the starting library to define a cell fitness score. Negative cell fitness scores indicate relative depletion whereas positive scores indicate relative enrichment of cells bearing these sgRNAs.

**Figure 1.**
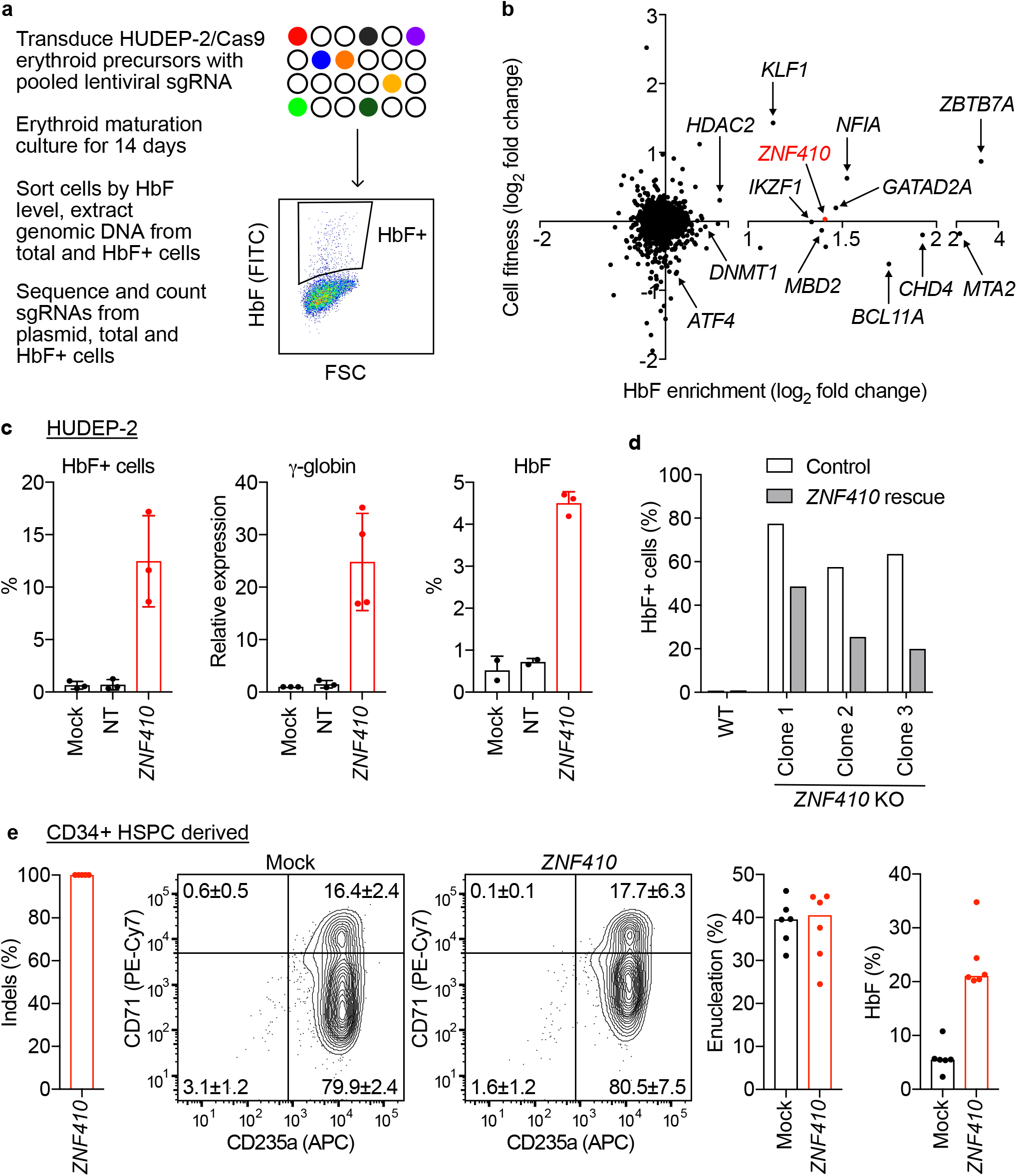
*ZNF410* is a novel HbF repressor. (a) Schematic of CRISPR/Cas9-based knockout screen in HUDEP-2 cells to identify novel repressors of HbF expression. (b) HbF enrichment and cell fitness scores for each of 1591 transcription factors and 13 genes of the NuRD complex. The gene *ZNF410* was prioritized for further study based on positive HbF enrichment score, neutral cell fitness score and unknown role in erythropoiesis and globin regulation. (c) HUDEP-2/Cas9 cells nontransduced (mock) or transduced with nontargeting (NT) or *ZNF410* targeting sgRNA assayed on day 9 of erythroid differentiation with intra-cellular staining (HbF+ cells), RT-qPCR (*HBG1/2* expression, fold change relative to mock) and HPLC (HbF level). Bars indicate mean values and error bars standard deviation (n=3), with p<0.05 for each comparison of NT to *ZNF410* edited. (d) Intra-cellular HbF staining of HUDEP-2 wild-type (wt) cells and three *ZNF410* knockout HUDEP-2 clones without or with (gray bars) re-expression of *ZNF410*. (e) *ZNF410* targeted by RNP electroporation of Cas9 and sgRNA in CD34+ HSPCs and subsequently differentiated to erythroid cells *in vitro*. Bars indicate median value, experiments performed in 4 individual donors including biological triplicate for donor 4 (total n=6 replicates). At the end of erythroid culture (day 18), erythroid maturation was assessed by surface expression of CD71 and CD235a and enucleation frequency by Hoechst staining. Representative FACS plots are shown. Quadrant values indicated are mean ± SD; t-test comparing *ZNF410* edited to mock for CD71+CD235a+ and CD71-CD235a+ quadrants did not show significant differences (p>0.05). HbF level measured by HPLC was increased in *ZNF410* edited primary erythroid cells compared to mock control cells (p<0.0001).

As expected, we found that known HbF regulators like *BCL11A* and *ZBTB7A* showed highly elevated HbF enrichment scores (**Fig. 1b**). For *BCL11A* we observed a modest negative fitness score, suggesting that loss of this gene had a modest negative impact on cell accumulation in vitro. For *ZBTB7A* we observed a positive fitness score, suggesting cells mutated at this gene accumulated in the population, consistent with its known requirement for terminal erythroid maturation^11^. In addition, we validated prior findings that a NuRD subcomplex including *CHD4, MTA2, GATAD2A, MBD2*, and *HDAC2* was required for HbF control^6^. Editing *CHD4* led to potent HbF induction but was associated with negative cell fitness.

We observed sgRNAs targeting *ZNF410* were associated with robust HbF induction (**Fig. 1b**). Unlike other regulators like *BCL11A, ZBTB7A*, and *CHD4*, we observed no fitness effects of targeting *ZNF410*. Relatively few prior studies have investigated *ZNF410*, encoding a zinc finger protein with a cluster of five C2H2 zinc fingers. It has not been previously implicated in globin gene regulation or erythropoiesis. One previous study indicated that over-expression of ZNF410 in human foreskin fibroblasts led to increased expression of matrix remodeling genes *MMP1, PAI2* and *MMP12* and ZNF410 sumoylation extended its half-life^12^. The biochemical functions and biological roles of endogenous ZNF410 remain largely unexplored. Therefore we focused on ZNF410 as a potentially novel regulator of HbF.

### Validation in HUDEP-2 cells and primary adult erythroid precursors

We first tested the role of ZNF410 in HbF repression by targeting it in HUDEP-2 cells with individual gRNAs. Upon editing in a bulk population of cells, we found induction of HbF, as measured by HbF+ cells by intracellular flow cytometry, *HBG1/2* expression by RT-qPCR, and HbF induction by HPLC (**Fig. 1c**). We generated 3 single cell derived HUDEP-2 *ZNF410* biallelic KO clones. In each clone, the fraction of HbF+ cells was elevated. Upon re-expression of ZNF410, HbF was partially silenced, consistent with a causal role of ZNF410 in repressing HbF (**Fig. 1d**).

We next examined the role of ZNF410 in HbF repression in primary erythroblasts derived from erythroid culture of adult mobilized peripheral blood CD34+ HSPCs. Using cells from 3 independent donors, we found that targeting *ZNF410* by 3xNLS-SpCas9:sgRNA RNP electroporation produced >99% indels with a +1 insertion allele (**Fig. 1e**). *ZNF410* targeted erythroblasts displayed normal erythroid maturation based on immunophenotype and enucleation, yet robust induction of HbF from a median of 5.5% in mock to 21.1% in *ZNF410* targeted samples (**Fig. 1e**, p<0.0001).

### ZNF410 is a DNA-binding protein with highly restricted chromatin occupancy

We performed dense mutagenesis of *ZNF410* to identify critical minimal sequences required for function. In this experimental design, heightened HbF enrichment scores indicate sequences where not only frameshift but also in-frame mutations may be associated with loss-of-function^6^. We observed heightened HbF enrichment scores especially when targeting sequences from exons 6-9 encoding the cluster of five C2H2 zinc fingers of *ZNF410* (**Fig. 2a**). This dependence on its putative DNA binding domain suggested that the DNA-binding function of ZNF410 might be important for its role in HbF repression.

**Figure 2.**
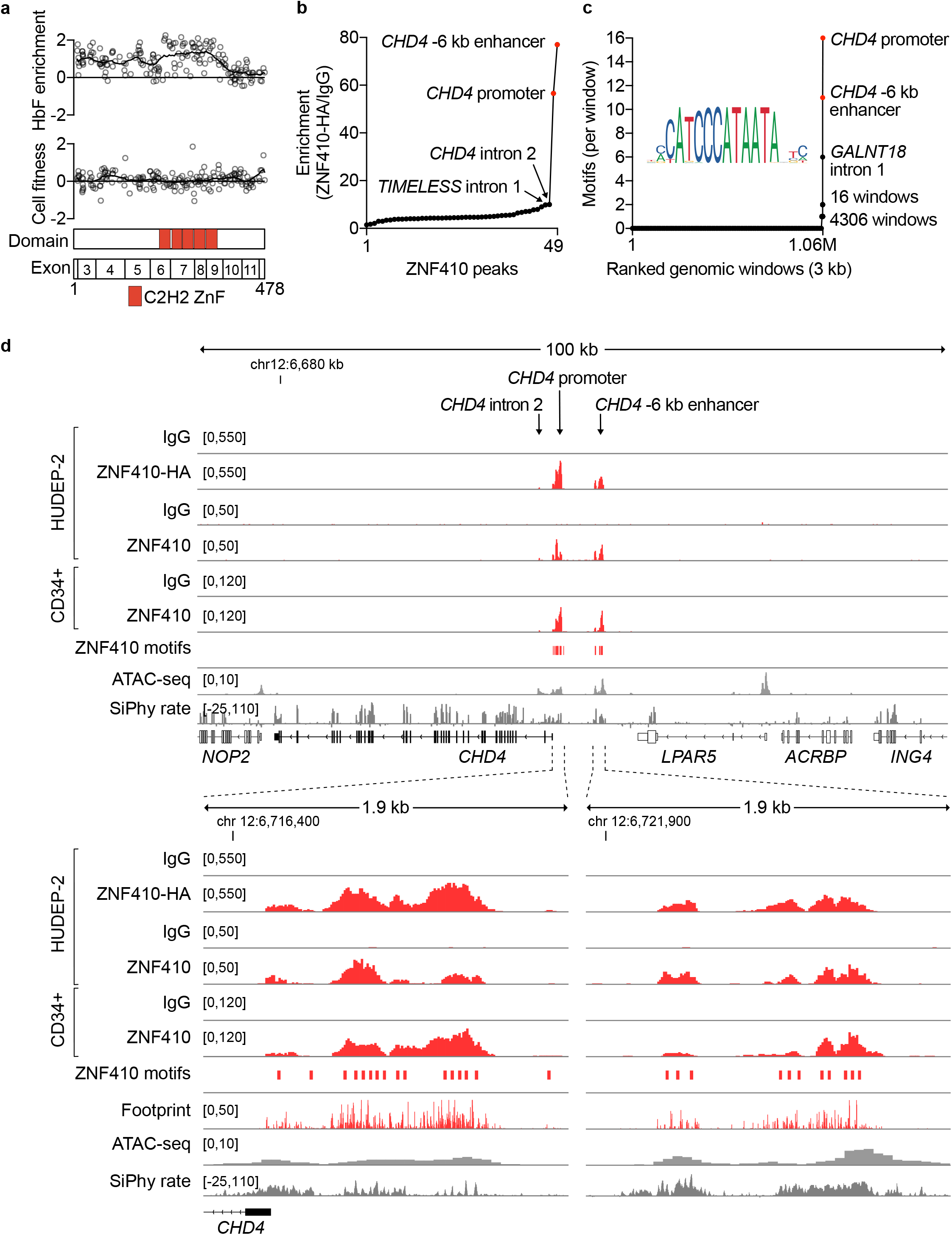
ZNF410 genomic chromatin occupancy is restricted to two *CHD4* elements with densely clustered motifs. (a) Dense mutagenesis of *ZNF410* coding sequence by pooled screening of 180 sgRNAs (NGG PAM restricted). Each circle represents enrichment score of an individual sgRNA, black line Loess regression. The 5 C2H2 zinc-finger domains (red rectangles) of *ZNF410* appear essential for HbF repression. (b) Genome-wide ZNF410 chromatin occupancy identified by CUT&RUN in HUDEP-2 samples with ZNF410-HA over-expression using anti-HA antibody compared to IgG control (n=4 for each). The two peaks with greatest enrichment of ZNF410 binding were at the *CHD4* promoter and *CHD4* −6 kb enhancer. The next most enriched peaks, at *CHD4* intron 2 and *TIMELESS* intron 1, showed substantially less enrichment. (c) Genome-wide ZNF410 motif occurrences (identified from JASPAR and mapped by pwmscan) across 3 kb sliding windows. Only three windows comprised more than two ZNF410 motifs, including the *CHD4* promoter (16 motifs), *CHD4* −6 kb enhancer (11 motifs), and *GALNT18* intron 1 (6 motifs). (d) *CHD4* locus at 100 kb (top panel) or 1.9 kb resolution (bottom panels) indicating ZNF410 binding (red peaks) at the *CHD4* promoter and *CHD4* −6 kb enhancer regions in representative control IgG (n=9) and anti-HA (n=7) samples in HUDEP-2 cells over-expressing HA-tagged ZNF410, control IgG (n=1) and anti-ZNF410 (n=1) in HUDEP-2 cells, and control IgG (n=2) and anti-ZNF410 (n=2) in CD34+ HSPC derived erythroid precursors. Positions of ZNF410 motifs (red rectangles), cleavage frequency (footprint) from ZNF410-HA CUT&RUN (red bars), accessible chromatin by ATAC-seq (gray peaks, n=3) and DNA sequence conservation by SiPhy rate.

We examined the chromatin occupancy of ZNF410 by conducting CUT&RUN, an approach to studying protein-DNA interactions in situ without fragmentation or cross-linking^13^. Initially we used an HA antibody to probe for epitope tagged ZNF410 in HUDEP-2 cells. Known HbF repressing TFs like BCL11A and ZBTB7A act by binding to proximal promoter elements at the fetal *γ*-globin (*HBG1* and *HBG2*) genes. We did not observe any chromatin occupancy of ZNF410 at the *α*-globin (*HBA1* and *HBA2*) or *β*-globin (*HBB*) gene clusters (**Supp. Fig. 1a, b**). Unlike typical DNA binding transcription factors which show thousands of binding sites genome wide, ZNF410 showed highly restricted chromatin occupancy. With standard peak calling parameters, we found 49 peaks, but most of these had marginal enrichment of ZNF410-HA signal compared to an IgG control. The top two peaks were found at the *CHD4* locus, one at the promoter (57-fold enrichment) and the other 6 kb upstream at a region of open chromatin (77-fold enrichment, **Fig. 2b, 2d**). This latter element we subsequently refer to as the *CHD4* −6 kb enhancer. The third most enriched peak was in *CHD4* intron 2, with ~10 fold enrichment (**Fig. 2b, 2d**). The fourth most enriched peak was in *TIMELESS* intron 1, with ~10 fold enrichment, around accessible chromatin at sequences bearing an LTR element (**Fig. 2b**, **Supp. Fig. 2a**).

Subsequently we used a ZNF410 antibody to probe for endogenous ZNF410 with CUT&RUN, both in HUDEP-2 cells and in CD34+ HSPC derived erythroid precursors (**Fig. 2d**). In both cases, we found that ZNF410 chromatin occupancy was highly restricted to *CHD4*. In HUDEP-2 cells, only 5 total peaks were identified, the top 4 of which were at the *CHD4* promoter and *CHD4* −6 kb enhancer (**Fig. 2d**, **Supp. Fig. 2b**). The 5th peak was at intronic sequences of *DPY19L3* bearing an LTR element (5-fold enrichment) (**Supp. Fig. 2d**). In CD34+ HSPC-derived erythroid precursors, only 5 total peaks were identified, all of which were at the *CHD4* promoter or *CHD4* −6 kb enhancer, with no other genomic sites of ZNF410 occupancy found (**Fig. 2d**, **Supp. Fig. 2c**).

The ZNF410 binding motif has previously been described by high-throughput SELEX using expression of the DNA-binding domain in 293FT cells^14^. We observed a striking cluster of these motifs at *CHD4*, with numerous motif instances found at both the promoter (16 motifs) and the −6 kb enhancer (11 motifs, **Fig. 2c, d**). We scanned the genome for the ZNF410 binding motif, dividing the genome into 3 kb windows. 4306 genomic windows had 1 motif instance and 16 windows had 2 motif instances (**Fig. 2c**). Only 3 windows had more than 2 motif instances, of which 2 were the aforementioned *CHD4* elements. We observed 6 motif instances within a window at *GALNT18* intron 1, although we observed neither ZNF410 occupancy nor chromatin accessibility at this locus in erythroid precursors (**Supp. Fig. 1c**).

### ZNF410 regulates HbF through CHD4

These results suggested that ZNF410 exhibits singular binding to *CHD4*. We performed RNA-seq of HUDEP-2 cells edited at *ZNF410* to measure gene expression changes (**Fig. 3a**). Based on log_2_ fold change >1 and adjusted p-value <0.01, there were 63 differentially expressed genes. *CHD4* was the most significantly downregulated gene upon *ZNF410* editing (L2FC −1.07, p_adj_ 2.27×10^-43^). *HBG2* was the 4th most significantly upregulated gene (L2FC 2.35, p_adj_ 5.93×10^-25^). Gene set enrichment analysis showed that genes differentially expressed after *ZNF410* editing were enriched in those differentially expressed after *CHD4* editing (for upregulated genes, NES 1.61, q 0.05; for downregulated genes, NES −1.39, q 0.09; **Fig. 3b, Supp. Fig. 3a**). The expression of *ZNF410* and *CHD4* were significantly correlated across 54 human tissues from the GTEx dataset^15^ (**Supp. Fig. 3b**, Pearson correlation, r 0.77, p<0.0001). We evaluated a repository of genome-wide CRISPR KO screen data spanning 558 cell lines to identify genes with a similar pattern of cellular dependency as *ZNF410*^16,17^. We found that *CHD4* was the most similarly codependent gene across cell lines, indicating a pervasive relationship between *ZNF410* and *CHD4* (**Fig. 3c**). These results suggest that a major function of ZNF410 across numerous cellular contexts appears to be control of CHD4 expression.

**Figure 3.**
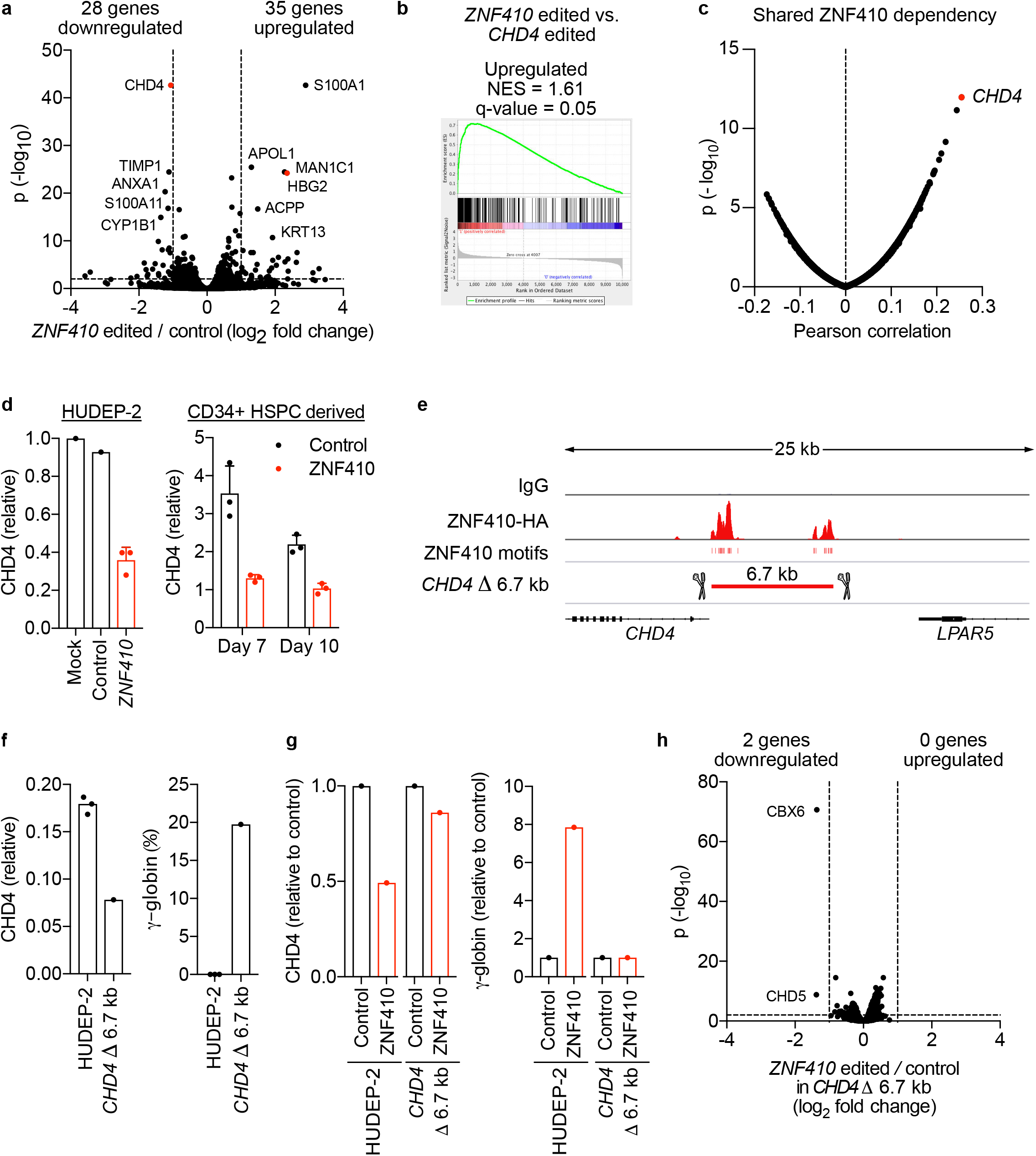
ZNF410 represses HbF by activating *CHD4*. (a) RNA-seq differential gene expression analysis of *ZNF410* (n=3) compared to *AAVS1* (n=3) targeted HUDEP-2 cells. Downregulated and upregulated genes defined by p_adj_<0.01 and L2FC<-1 or >1 respectively. (b) Comparison of genes upregulated in *ZNF410* and *CHD4* mutant cells by GSEA shows enrichment of *CHD4* regulated genes in the *ZNF410* regulated gene set. (c) Pearson correlation between *ZNF410* dependency and *CHD4* dependency across 558 cell lines identifies *CHD4* as the most *ZNF410* codependent gene. (d) *CHD4* expression measured by RT-qPCR in *ZNF410* targeted (n=3) compared to mock and *AAVS1* targeted control HUDEP-2 cells (left panel) and in *ZNF410* targeted primary erythroblasts derived from CD34+ HSPCs (n=3, p<0.01) compared to safe sgRNA targeted control cells on day 7 and day 10 of erythroid culture (right panel). (e) Cas9 paired cleavages with CHD4-proximal-gRNA-1 and CHD4-distal-gRNA-1 were used to generate an element deletion clone *(CHD4* Δ 6.7 kb), with the biallelic deletion spanning both of the ZNF410 binding regions upstream of *CHD4*. (f) *CHD4* expression measured by RT-qPCR in the *CHD4* Δ 6.7 kb clone compared to 3 individual HUDEP-2 cell clones plated in parallel. *HBG* expression relative to total *β*-like globin (*HBG*+*HBB*) measured by RT-qPCR in the *CHD4* Δ 6.7 kb deletion clone compared to control clones. (g) CHD4 Δ 6.7 kb clones and HUDEP-2 cells were subjected to control (safe) and *ZNF410* targeting by RNP electroporation. Relative *CHD4* and *HBG* expression measured by RT-qPCR. (h) RNA-seq differential gene expression analysis of *ZNF410* targeted (n=3) compared to *AAVS1* targeted (n=3) *CHD4 Δ* 6.7 kb clones. Downregulated and upregulated genes defined by p_adj_<0.01 and L2FC<-1 or >1 respectively.

We validated the changes in CHD4 expression after *ZNF410* editing by RT-qPCR in both HUDEP-2 cells and primary erythroid precursors derived from CD34+ HSPCs. We found that CHD4 mRNA expression was reduced by 57% after *ZNF410* editing (**Fig. 3d**, p<0.01). To test the requirement of ZNF410 binding for CHD4 expression, we generated HUDEP-2 cell clones in which the two upstream ZNF410 motif clusters at *CHD4* were both deleted by paired genomic cleavages (**Fig. 3e, Supp. Fig. 3c**). We isolated 4 biallelically deleted HUDEP-2 clones. We found that CHD4 expression decreased by 56-79% after deletion of the upstream elements, similar to the decrease observed after editing *ZNF410* itself. Consistent with reduced expression of CHD4, *γ*-globin was induced (**Fig. 3f, Supp. Fig 3d, 3e**). No change in CHD4 expression was observed upon *ZNF410* editing in the absence of the upstream elements, suggesting that the control of CHD4 expression requires these elements. We did not observe further *γ*-globin induction in *CHD4* upstream element deleted cells upon *ZNF410* editing (**Fig. 3g, Supp. Fig. 3e**). In contrast, *γ*-globin increased in these same cells upon *ZBTB7A* editing, indicating the cells were competent for further *γ*-globin induction. We performed RNA-seq of *CHD4* Δ6.7 kb element deleted cells after *ZNF410* editing (**Fig. 3h**). In contrast to HUDEP-2 cells, we only observed 2 differentially expressed genes after *ZNF410* editing in *CHD4* Δ6.7 kb element deletion cells, consistent with our prior results that nearly all gene expression changes found after *ZNF410* editing are due to changes in CHD4 expression. Together these results suggest that ZNF410 represses *γ*-globin exclusively by binding upstream elements and trans-activating CHD4.

### ZNF410 is a non-essential gene

ZNF410 and its mouse ortholog Zfp410 share 94% amino acid identity, including 98% at the cluster of 5 ZnFs^12^. We performed CUT&RUN to investigate the chromatin occupancy of endogenous Zfp410 in a mouse erythroid cell line (MEL cells). Similar to results in human erythroid precursors, we observed that genomic enrichment of Zfp410 binding was highly restricted to the *Chd4* locus, with 77-fold enrichment at the promoter and 45-fold enrichment at the *Chd4* −6 kb enhancer, each overlapping accessible chromatin regions and motif clusters (**Fig. 4a, b**). The third most enriched site for Zfp410 occupancy was at the promoter of *Hist1h2bl*, with ~14 fold enrichment, although no motifs were observed at this site (**Supp. Fig. 4a**). To evaluate the requirement of ZNF410 in normal development and homeostasis, we investigated mice with a loss-of-function allele of the mouse ortholog *Zfp410*. We obtained mouse embryonic stem cells that are heterozygous for a *Zfp410* gene trap allele (Gt) from the European Mouse Mutant Cell Repository (EuMMCR). The targeting cassette was inserted to intron 5 to disrupt expression of full-length Zfp410 (**Supp. Fig. 4b**). Of note, exons 6-9 encode the five ZnFs (**Supp. Fig. 4c**). We derived heterozygous mice with germline transmission of this allele. Although the sample size is currently small, we observed 6 *Zfp410^Gt/Gt^* homozygotes out of 20 live births from Zfp410^+/Gt^ heterozygote intercrosses, consistent with expected Mendelian transmission (**Fig. 4c**). *Zfp410* expression was reduced by >98% in *Zfp410^Gt/Gt^* homozygous mouse whole blood (**Fig. 4d**). The *Zfp410^Gt/Gt^* homozygotes showed moderately reduced body weight compared to heterozygotes or wt mice (**Fig. 4e**), but otherwise appeared healthy and active. Analysis of complete blood counts showed apparently unremarkable hematologic parameters in *Zfp410^Gt/Gt^* homozygous mice, including no evidence of anemia or hemolysis (**Fig. 4f**). The absence of a severe phenotype of constitutive *Zpf410* loss-of-function is notable in comparison to other HbF regulators. For example, *Bcl11a* deficient mice experience perinatal lethality^18^, *Zbtb7a* deficient mice mid-gestation embryonic lethality due to anemia^11^, and *Chd4* deficient mice pre-implantation embryonic lethality^19^. Together these results suggest that ZNF410 is an evolutionarily conserved HbF repressor that is not essential for vertebrate survival.

**Figure 4.**
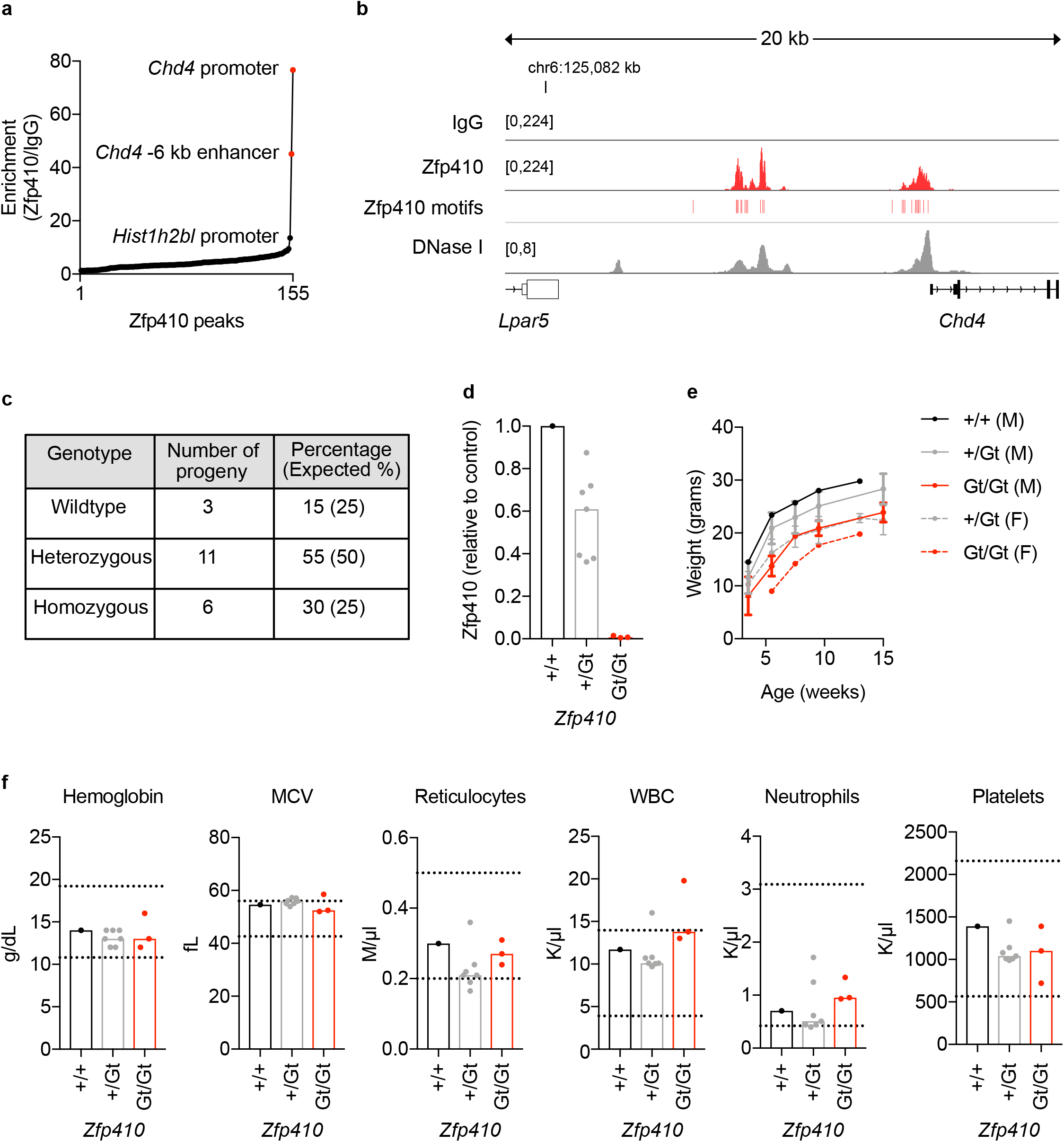
*Zfp410* deficient mice are viable with unremarkable hematology. (a) CUT&RUN in mouse erythroleukemia (MEL) cells using anti-Zfp410 antibody (n=3) and IgG control (n=3). Enrichment for Zfp410 binding concentrated at *Chd4* promoter (~77 fold enrichment) and *Chd4* −6 kb enhancer (~45 fold enrichment) peaks. The next most enriched peak was at the *Hist1h2bl* promoter (~14 fold enrichment). (b) *Chd4* locus showing Zfp410 binding (red peaks) at the *Chd4* promoter and *Chd4* −6 kb enhancer in representative IgG control (n=3) and anti-Zfp410 (n=3) samples. Positions of Zfp410 motifs (red rectangles) and accessible chromatin by DNase-seq (gray peaks). (c) Mouse ES cells heterozygous for *Zfp410* gene-trap allele (Gt), obtained from EuMMCR, were used to generate heterozygous (*Zfp410* +/Gt) and homozygous (*Zfp410* Gt/Gt) gene-trap mice, with *Zfp410* +/Gt intercrosses yielding 20 progeny from 4 litters. (d) *Zfp410* expression, measured by RT-qPCR using primers spanning exons 5 and 6, was diminished in *Zfp410* Gt/Gt (n=3) mouse peripheral blood compared to heterozygous (n=7, p<0.05) and wildtype control animals. (e) Mouse weight was measured at indicated time points over the course of 15 weeks. (f) Peripheral blood hematological parameters, with normal ranges for hemoglobin, mean corpuscular volume (MCV), reticulocyte, white blood cell (WBC), neutrophil and platelet count shown by dotted lines.

### ZNF410 appears dispensable for human erythropoiesis and hematopoiesis

To evaluate the role of *ZNF410* in human hematopoiesis, we performed gene editing of *ZNF410* in primary human hematopoietic stem and progenitor cells (HSPCs). We electroporated 3xNLS-SpCas9 and sgRNA as ribonucleoprotein (RNP) to CD34+ HSPCs from two healthy donors and achieved >99% indels (**Fig. 5a, b**). Since all of these measured indels were +1 insertions, biallelic *ZNF410* knockouts comprised nearly all cells in the population. To test the role of *ZNF410* more broadly in hematopoiesis, we performed xenotransplantation of edited HSPCs to immunodeficient NBSGW mice (**Fig. 5a**). NBSGW mice support multilineage (lymphoid, myeloid and erythroid) human engraftment in absence of conditioning therapy^20^. After 16 weeks we analyzed bone marrow from engrafted recipients. We observed similar human hematopoietic engraftment for *ZNF410* edited HSPCs compared to mock control xenografts (**Fig. 5c**). *ZNF410* +1 insertion frameshift indels were observed at >99% frequency in total BM human hematopoietic cells similar to the input cell product (**Fig. 5b**). A comparable distribution of multilineage hematopoietic reconstitution was found in control and *ZNF410* edited recipients, including B-lymphocyte, T-lymphocyte, granulocyte, monocyte, HSPC and erythroid contributions (**Fig. 5d, e**). We found that CHD4 expression was decreased by ~60% in human erythroid cells sorted from bone marrow, similar to *in vitro* results (**Fig. 5f**). The level of HbF as measured by HPLC from engrafting human erythrocytes was ~2.5% in controls and ~17% in *ZNF410* edited recipients (**Fig. 5g**).

**Figure 5.**
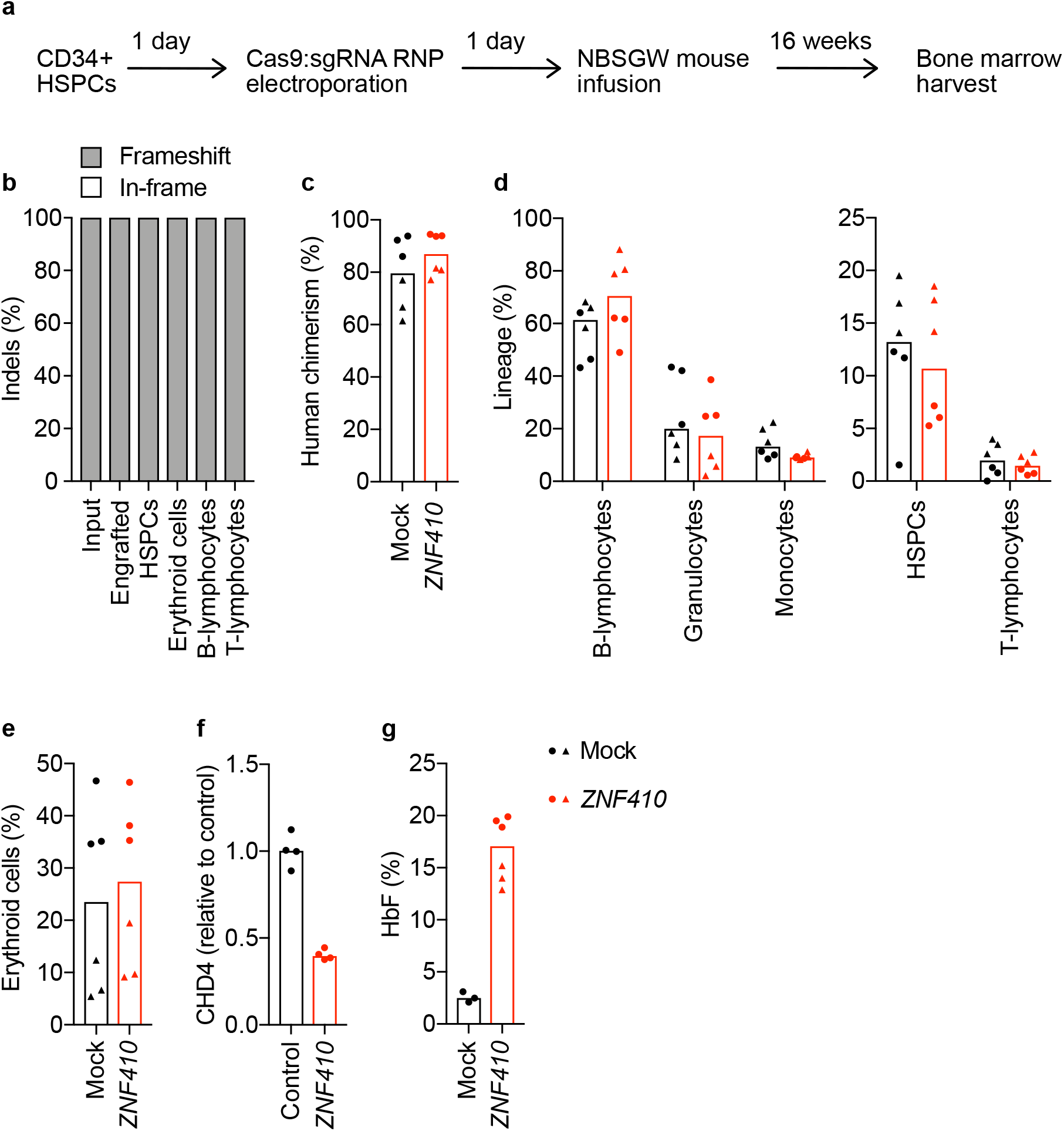
*ZNF410* deficient human HSPCs de-repress HbF and retain repopulation potential. (a) Schematic of gene editing and transplant of human CD34+ HSPCs in immunodeficient NBSGW mice. Animals were euthanized 16 weeks post-transplant and bone marrow (BM) was harvested and sorted into various subpopulations by flow cytometry. (b-e) Two independent CD34+ HSPC donors were edited and transplanted into 6 mice for each condition (mock or *ZNF410* edited). Each symbol represents one mouse, recipients of donor 1 depicted as circles and donor 2 as triangles. Bars indicate median value. (b) Indel frequency at *ZNF410* was quantified in input cells 4 days after electroporation and in total and sorted engrafted BM cells. Percentage of frameshift alleles is represented in gray and the percentage of in-frame alleles is represented in white for each bar. (c) Engraftment of human hematopoietic cells assessed by hCD45+ compared to total CD45+ cells. (d) B-lymphocytes (CD19+), granulocytes (CD33-dim SSC-high) and monocytes (CD33-bright SSC-low) expressed as fraction of hCD45+ cells. HSPCs (CD34+) and T-lymphocytes (CD3+) expressed as fraction of hCD45+ CD19-CD33-cells. (e) Erythroid cells (hCD235a+) expressed as fraction of hCD45-mCD45-cells. (f) *CHD4* expression measured by RT-qPCR in human erythroid cells from control (n=4) and *ZNF410* edited (n=4) xenografts. (g) HbF measured by HPLC from hemolysates of sorted BM hCD235a+ cells.

For comparison, we also performed xenotransplant experiments with *BCL11A* and *ZBTB7A* edited HSPCs. Consistent with the known role of *BCL11A* in supporting HSC self-renewal (and unlike the selective erythroid impact of *BCL11A* erythroid enhancer editing^21^), we observed reduced human chimerism in the bone marrow of recipients of *BCL11A* exon edited HSPCs after 16 weeks, reduced *BCL11A* edits compared to input cell product, and reduced fraction of frameshift alleles compared to total edits (**Supp. Fig. 5a, b**). For *ZBTB7A*, the fraction of engrafting human hematopoietic cells was similar to controls but the gene edits were reduced compared to input cell product (**Supp. Fig. 5a, b**). During erythroid maturation culture, *ZBTB7A* edited HSPCs showed impaired terminal erythroid maturation potential based on immunophenotype and enucleation frequency, in contrast to *ZNF410* edited cells (**Supp. Fig. 5c, d**). Together these results suggest HSPCs bearing *BCL11A* and *ZBTB7A* loss-of-function alleles are under negative selective pressure during hematopoietic repopulation and erythropoiesis unlike *ZNF410* edited cells.

## Discussion

The advances in knowledge of the molecular details of hemoglobin switching have begun to bear fruits in the form of novel autologous therapies^1^. A host of HSC-based therapies that reduce the expression of BCL11A in erythroid cells or prevent its binding to *HBG1/2* promoter sequences are in clinical trials or late preclinical development. However the clinical unmet need remains vast, with ~300,000 infants estimated to be born each year worldwide with sickle cell disease and tens of thousands more with severe *β*-thalassemia. The feasibility in terms of cost and infrastructure to scale up autologous cell-based therapies remains uncertain. Furthermore the toxicity of myeloablative transplantation will likely render these therapies out of reach to many patients.

The most realistic near-term hope to develop scalable therapies to address the root cause of these diseases would be through pharmacotherapy. Drugs that could interrupt molecular vulnerabilities required for adult erythroid cells to maintain fetal globins in a silenced state are greatly needed. These could complement or even supplant existing treatments like hydroxyurea^22^. BCL11A itself would certainly represent a preeminent target. Its roles in erythropoiesis besides HbF silencing are modest. However BCL11A plays essential roles in various hematopoietic lineages, including in B-lymphocytes, dendritic cells and hematopoietic stem cells^18,23–25^. In addition, it has functions beyond hematopoiesis not only in the central nervous system but also in breast and pancreatic cells^26,27^. Another exciting target would be ZBTB7A given its potent role in HbF repression. However ZBTB7A is required for terminal erythropoiesis and germinal center B cell maturation and plays important roles in T-lymphocytes, osteoclasts and HSCs^28^. A specific NuRD subcomplex including CHD4, GATAD2A, MBD2, MTA2 and HDAC2 is required for HbF silencing^6,29^. Targeting NuRD including key protein-protein interactions appears promising but would need to navigate the numerous gene expression programs that depend on this chromatin complex. For most of the known HbF regulators, their pleiotropic roles could yield potential on-target liabilities with narrow therapeutic index even if rational targeting approaches could be devised.

Here we identify ZNF410 as a novel HbF repressor that acts specifically to enhance the expression of CHD4. Complete knockout of ZNF410 is well-tolerated, apparently since the remaining level of CHD4 is sufficient to maintain cellular functions. *Zfp410* mutant mice survive to adulthood and *ZNF410* knockout HSPCs demonstrate no defects in erythroid maturation or hematopoietic reconstitution. Traditionally TFs have been considered undruggable targets. However the example of small molecules binding and resulting in specific degradation of zinc finger proteins like IKZF1 has encouraged the development of ligands to modulate DNA-binding factors^30,31^.

ZNF410 appears to represent a special form of gene regulator. Conventional DNA-binding TFs bind and directly control the expression of thousands of genomic targets. In contrast, ZNF410 shows unique binding to *CHD4*. This exquisite specificity appears to be achieved through a remarkable clustering of 27 ZNF410 binding sites at the *CHD4* promoter and −6 kb enhancer, a density unlike anywhere else in the genome. Both *ZNF410* itself and its two target elements at *CHD4* are highly conserved across vertebrates. Despite thousands of ZNF410 motifs across the genome, we detected minimal ZNF410 occupancy at these sites. The absence of detectable ZNF410 occupancy or chromatin accessibility even at *GALNT18* intron 1 with 6 clustered motif instances suggests that motif clusters may be necessary but insufficient for ZNF410 binding. Another example of clustered homotypic TF binding sites associated with gene control is the binding of ZFP64 to the *MLL* gene promoter, activating the expression of the chromatin regulator MLL, although in this case ZFP64 shows a limited set of additional direct target genes^32^. CHD4 is an especially abundant nuclear protein in erythroid precursors^33^.

Haploinsufficiency of *CHD4* (or *MLL*) causes impaired intellectual development and congenital anomalies, suggesting that chromatin regulatory complexes must be maintained at precise levels to maintain proper gene regulation, particularly during development^34–36^. There are more than a thousand putative DNA-binding TFs, for many of which the genomic binding sites and regulons remain poorly characterized^37^. ZNF410 may be emblematic of a class of TFs relying on homotypic motif clusters^38^ with limited gene targets that are devoted to maintenance of core cellular programs.

In summary, here we identify ZNF410 as a dispensable TF that represses HbF level in adult-stage erythroid precursors by devoted maintenance of NuRD subcomplex levels through binding a singular cluster of sequences upstream of *CHD4*.

## Methods

### Cell culture

HUDEP-2 cells^39^ were cultured as previously described^40^. Expansion phase medium for HUDEP-2 cells consists of SFEM (Stemcell Technologies #09650) base medium supplemented with 50 ng/ml recombinant human SCF (R&D systems #255-SC), 1 μg/ml doxycycline (Sigma Aldrich #D9891), 0.4 μg/ml dexamethasone (Sigma Aldrich #D4902), 3 IU/ml EPO (Epoetin Alfa, Epogen, Amgen) and 2% Penicillin-Streptomycin solution (10,000 U/mL stock). Erythroid differentiation medium (EDM) for HUDEP-2 cells consists of Iscove’s Modified Dulbecco’s Medium (IMDM, ThermoFisher #12440053) supplemented with 1% L-Glutamine (Gibco #25030081), 330 μg/mL human holo-Transferrin (Sigma #T0665), 10 μg/mL human insulin (Sigma #I9278), 2 IU/mL heparin (Sigma #H3149), 5% inactivated human plasma (Octaplas, blood group AB, Octapharma), 3 IU/mL EPO (Epoetin Alfa, Epogen, Amgen) and 2% Penicillin-Streptomycin solution (10,000 U/mL stock). EDM-2 medium for HUDEP-2 cells is EDM supplemented with 100 ng/ml human SCF and 1 μg/ml doxycycline. CD34+ HSPCs from adult mobilized peripheral blood from de-identified healthy donors were purchased from Fred Hutchinson Cancer Research Center, Seattle, Washington. Upon thawing, CD34+ HSPCs were resuspended in X-VIVO 15 medium (Lonza #04-380Q) containing 50 ng/ml recombinant human Flt-3 ligand (Peprotech #300-19), 100 ng/ml recombinant human TPO (Peprotech #300-18) and 100 ng/ml recombinant human SCF (R&D systems #255-SC) (referred to as X-VIVO complete medium). Erythroid differentiation was performed in 3 phases as previously described^41^. Mouse erythroleukemia cells, MEL-745A cl. DS19, were cultured in RPMI 1640 medium supplemented with 10% fetal bovine serum and 1% Penicillin-Streptomycin.

### sgRNA library screening

For library screening, HUDEP-2 cells with stable expression of LentiCas9-Blast (Addgene plasmid 52962) were transduced at a low multiplicity of infection (MOI) with virus containing sgRNA library cloned in lentiGuide-Puro (Addgene plasmid 52963) to ensure that most cells received only one sgRNA in expansion phase medium^42^. The sgRNA library included 18,020 gRNA overlapping those in GeCKOv2^43^ and Avana^44^ libraries targeting 1591 transcription factors and 13 genes of the NuRD complex as controls. After 24 hours, cells were transferred to and cultured in erythroid differentiation medium for 14 days. At the end of erythroid culture, cells were processed for intra-cellular HbF staining using Fetal Hemoglobin Monoclonal Antibody (HBF-1) conjugated to FITC (Thermo Fisher #MHFH01), and HbF+ cells were sorted by FACS as previously described^6^. Genomic DNA was extracted from the total cell population and from HbF+ sorted cells and deep sequenced to identify guide RNAs with enrichment in the HbF+ pool as previously described^6^. Briefly, two step PCR was performed to amplify sgRNA cassette from genomic DNA, using Herculase II Fusion DNA polymerase (Agilent #600677). Multiple reactions of the first PCR were set up for each sample in order to maximize genomic DNA input up to 1000 cell equivalents per sgRNA. After the first PCR, all reactions for each sample were pooled and 1 ul of this mix used as input for the second PCR reaction which was performed in duplicate. Illumina adaptor and sample barcodes added in the second PCR. Primers for the second PCR were of variable length to increase library complexity^42^. Sequences of PCR primers can be found in the **Supplementary Table**. Amplicons obtained from the second PCR were purified by gel extraction and quantified using the Qubit dsDNA HS assay kit (Invitrogen #Q32851). Single-end 75 bp sequencing was performed on the NextSeq 500 platform by the Molecular Biology Core Facilities at Dana-Farber Cancer Institute. Candidate HbF regulators were identified by analyzing sequencing data using the model-based analysis of genome-wide CRISPR-Cas9 Knockout (MAGeCK) computational tool^45^.

### Validation in HUDEP-2 cells

Candidate HbF regulators identified by the screen were validated in arrayed format in HUDEP-2 cells. HUDEP-2 cells with stable expression of lentiCas9-Blast were transduced with sgRNA cloned in lentiGuide-Puro in expansion phase medium. 24 hours after transduction, cells were cultured in EDM2 for 4 days, EDM with doxycycline for 3 days and EDM without doxycycline for 2 days as previously described^40^. These culture conditions result in differentiation of normal HUDEP-2 cells to orthochromatic erythroblasts. At the end of erythroid differentiation cells were divided into aliquots and processed for intra-cellular HbF staining, RNA isolation and hemoglobin HPLC. In addition to mock treated cells, non-targeting sgRNAs or sgRNAs targeting either *AAVS1* or a functionally neutral locus on chr2 (so-called “safe targeting” sgRNA)^46^ were used as experimental controls as indicated in each figure legend. For RNA sequencing experiments HUDEP-2 cells were cultured in expansion phase medium for 6 days after transduction or electroporation. RNA was isolated using Trizol according to the manufacturer’s protocol (Thermo Fisher #15596026). Purified RNA was treated with DNase I. mRNA libraries were prepared and sequenced by the Molecular Biology Core Facilities at Dana-Farber Cancer Institute.

### Generation of *ZNF410* null HUDEP-2 cell clones

The entire coding sequence of ZNF410 was deleted in HUDEP-2 cells using paired Cas9 cleavages. *ZNF410* null HUDEP-2 cell clones were generated in two steps. In the first step a cell clone with heterozygous deletion of *ZNF410* was obtained using the gRNAs ZNF410-del-5’-tgt1 and ZNF410-del-3’-tgt1. In the second step this heterozygous *ZNF410* null clone was retargeted using a second pair of guide RNAs, ZNF410-del-5’-tgt2 and ZNF410-del-3’-tgt2, to obtain biallelic deletion of ZNF410. Three individual *ZNF410* null clones were obtained by limiting dilution of bulk edited cells. Mono- or biallelic deletion clones were identified by PCR amplification of the genomic DNA flanking the deletion (outer PCR) and inside the targeted region (inner PCR) using the following primers: ZNF410-outer-FP/RP and ZNF410-inner-FP/RP. For the rescue experiment, the three *ZNF410* null clones were transduced with either an HA-tagged *ZNF410* construct or an HA-tagged nuclear localization sequence (NLS) containing control vector. Successfully transduced cells were obtained by selection of cells using blasticidin (Invivogen #ant-bl-05).

### Validation in CD34+ HSPCs

CD34+ cells were thawed and maintained in X-VIVO complete medium for 24 hours. 100,000 cells per condition were electroporated using the Lonza 4D nucleofector with 100 pmols 3xNLS-Cas9 protein and 300 pmols modified sgRNA targeting the gene of interest. In addition to mock treated cells, *AAVS1* targeting or “safe-targeting” RNPs were used as experimental controls as indicated in each figure legend. After electroporation cells were differentiated to erythroblasts as described previously^41^. 4 days after electroporation, genomic DNA was isolated from an aliquot of cells, the sgRNA targeted locus was amplified by PCR and processed for Sanger sequencing. Sequencing results were analyzed by Synthego’s ICE algorithm to obtain editing efficiency and allele contributions. At the end of erythroid culture (day 18) cells were processed for surface marker / enucleation analysis by staining with anti-CD71 (PE-Cy7 conjugated, eBioscience #25-0719-42), anti-CD235a (APC conjugated, eBioscience #17-9987-42) and Hoechst 33342 (Invitrogen #H3570) following manufacturer’s recommendations for antibody concentration and flow cytometry data acquisition on the BD LSR Fortessa. Cells were also processed for hemoglobin HPLC using the Bio-Rad D-10 hemoglobin testing system.

### Dense mutagenesis of *ZNF410*

180 guide RNAs were identified by searching for 20-mer sequences upstream of an NGG PAM on the sense and antisense strands of the consensus coding sequence (CCDS) for ZNF410 obtained from the Ensembl genome browser (Transcript ID ENST00000555044.6). Lentiviral sgRNA libraries were synthesized as previously described^47^ and pooled screening was performed as described in the sgRNA library screening section above. Sequencing results were analyzed by the CRISPRO tool^48^. For each gRNA an HbF enrichment score was calculated comparing the abundance of the gRNA in HbF-high cells to the total cell pool at the end of erythroid culture. Cell fitness scores were calculated by comparing the abundance of the gRNA in cells at the end of erythroid culture to the starting library. The CRISPRO algorithm maps the cell fitness and HbF enrichment score to gene, transcript and protein coordinates and lists associated protein structural domains.

### CUT&RUN

CUT&RUN was performed to identify the genome wide ZNF410 / Zfp410 DNA binding profile as previously described^13^. The antibodies used were anti-HA antibody (ThermoFisher #71-5500) in HUDEP-2 cells expressing an HA-tagged ZNF410 construct or anti-ZNF410 (Abcam #ab174204) to detect endogenously expressed ZNF410 in HUDEP-2 and primary human CD34+ cells as well as endogenously expressed Zfp410 in MEL cells. Normal rabbit IgG polyclonal antibody (Millipore Sigma #12370) was used as a control for non-specific sequence enrichment. Anti-H3K27me3 antibody (Cell signaling Technology #9733) was used as a positive control for the steps leading up to the chromatin release. Protein A-MNase was kindly provided by Dr. Steve Henikoff. Sequencing libraries were prepared using the NEBNext Ultra™ II DNA Library Prep Kit for Illumina as previously described^9^. Paired-end 42bp sequencing was performed on the NextSeq 500 platform by the Molecular Biology Core Facilities at Dana-Farber Cancer Institute. Sequencing data analyses was adapted from previous protocols^9,13^. FastQC (Babraham Institute) was performed for all samples to check sequencing quality. Adapter sequences were trimmed with Trimmomatic^49^ with the following settings: “ILLUMINACLIP:$TRIMMOMATIC/adapters/TruSeq3-PE.fa:2:15:4:4:true SLIDINGWINDOW:4:15 MINLEN:25.” Trimmed reads were aligned to the human reference genome hg19 using bowtie2^50^ with the following settings: “--end-to-end --no-unal --no-mixed --no-discordant --dovetail --phred33 -p 4.” The resulting alignment files (.sam) were converted to sorted, indexed bam files and marked for duplicates using Picard (https://broadinstitute.github.io/picard/). Reads were filtered using an alignment score cutoff of 10 with samtools^51^. Peak calling was performed using macs2^52^ with the following settings: “callpeak -f BAMPE -t [test replicates] -c [control replicates] -B -g [hs or mm] -q 0.05 -n [outputID].” Genomic regions annotated as part of the ENCODE project blacklist^53^ as problematic regions for alignment of high-throughput functional genomics data were excluded from analysis using files ENCFF001TDO (hg19, Birney lab, EBI) and ENCFF547MET (mm10, Kundaje lab, Stanford) and BEDtools^54^. Locus footprinting was performed to identify regions of DNA that are relatively protected from MNase cleavage compared to neighboring regions due to occupancy by a transcription factor. Footprint patterns at a locus were determined by enumerating the ends of each fragment sequenced and aligned to the locus. Data was visualized using IGV^55^.

### Genome-wide motif mapping

Genome-wide ZNF410 DNA binding motif instances were mapped using the pwmscan webtool (https://ccg.epfl.ch/pwmtools/pwmscan.php) and the ZNF410 motif MA0752.1 from the JASPAR CORE 2018 vertebrates motif library. The number of motif instances in the genome was enumerated using a 3 kb sliding window with a 100 bp overlap to determine genomic distribution of motif occurrence for ZNF410. Motifs that fall within the overlapping region between genomic windows are assigned to the adjacent window with the greater number of motifs, or if both adjacent windows have the same number of motifs, motifs are assigned to the first of the two windows.

### ATAC-seq and DNase-seq identification of regions of open chromatin

ATAC-seq was performed in HUDEP-2 cells grown in expansion phase medium following the OMNI-ATAC protocol^56^. MEL DNase-seq data was obtained from the ENCODE project^57,58^ (https://www.encodeproject.org/) from the lab of John Stamatoyannopoulos, UW (dataset: ENCSR000CNN, file: ENCFF990ATO).

### DNA sequence conservation

SiPhy rate^59^ (10 mer) from: http://www.broadinstitute.org/igvdata/hg19/omega.10mers.wig.tdf.

### GSEA

Genes that were differentially expressed in *ZNF410* targeted HUDEP-2 cells compared to *AAVS1* targeted control cells were compared by gene set enrichment analysis (GSEA)^60,61^ to genes that were differentially expressed in *CHD4* targeted HUDEP-2 cells compared to non-targeting control cells^6^. The list of genes differentially expressed in *CHD4* targeted HUDEP-2 cells are genes that were differentially expressed in both datasets (q<0.05) when either the helicase domain or the CHDCT2 domain of *CHD4* were perturbed using sgRNA GGTGTCAGTGCCCTGAGCCC or GAATTCGGGCAATGGTAGCT respectively from previously published data^6^. The motivation for combining these two datasets is based on the observation that helicase domain targeting is toxic to cells while CHDCT2 domain targeting is better tolerated and so the combined list of differentially expressed genes better represents gene expression changes due to *CHD4* ablation than either dataset alone.

### Gene dependency correlation

Gene dependency scores for 558 cell lines were obtained from the Achilles Avana 20Q2 Public CERES dataset of the Depmap portal (DepMap, Broad (2020): DepMap 20Q2 Public. figshare. Dataset. https://doi.org/10.6084/m9.figshare.12280541.v4.)^16,17^. Project Achilles performs genome scale CRISPR/Cas9 loss of function screening in cancer cell lines and uses CERES to determine a dependency score for each gene in each cell line. Pearson’s correlations of dependency scores and p-values were calculated for ZNF410 and every other gene in the dataset.

### Analysis of gene expression across human tissues

*ZNF410* and *CHD4* expression values (TPM) across 54 human tissues were obtained from the GTEx Portal^15^ on 10/01/2019. Pearson correlation was used to compare the expression of *ZNF410* and *CHD4*.

### Generation of *CHD4* Δ6.7 kb and Δ6.9 kb clones

The genomic region upstream of *CHD4* encompassing the two clusters of ZNF410 DNA binding motifs was targeted using a pair of sgRNAs (CHD4-proximal-gRNA-1: GUGCGGUGGGAUUUCCCGGC and CHD4-distal-gRNA-1: CGAGGCUGUGUCAGCGCCGC or CHD4-distal-gRNA-2: UUGGUCUGUGGGAUGGACAU) to generate HUDEP-2 clones with biallelic deletion of the intervening sequence. These clones are termed CHD4 Δ 6.7 kb (for clones generated using CHD4-proximal-gRNA-1 and CHD4-distal-gRNA-1) or CHD4 Δ6.9 kb (for clones generated using CHD4-proximal-gRNA-1 and CHD4-distal-gRNA-2). The bulk population of targeted cells was serial diluted and ~30 cells per plate were plated in 96 well plates to obtain single cell clones. Mono- or biallelic deletion was identified by PCR amplification of the genomic DNA flanking the deletion (outer PCR) and inside the targeted region (inner PCR) using the following primers: CHD4-Outer-FP and CHD4-Outer-RP1 or CHD4-Outer-RP2, CHD4-Inner-FP and CHD4-Inner-RP (sequences listed in **Supplementary Table**).

### RT-qPCR

RNA was isolated using either Trizol (Invitrogen #15596026) or the RNeasy Plus Mini kit (Qiagen #74136) following the manufacturer’s protocol. RNA was quantified using the Nanodrop spectrophotometer. cDNA was synthesized using the iScript cDNA synthesis kit (Bio-Rad #1708891) following the manufacturer’s recommendations. qPCR was performed using the Sybr Select Master Mix (Thermo Fisher #4472908) on an Applied Biosystems 7300 or Quant Studio 3 real-time PCR system. Primers used for RT-qPCR are listed in the **Supplementary Table**. *CAT* was used as a reference gene for human cells and *Gapdh* for mouse cells.

### Generation of *Zfp410* gene-trap allele mice

All animal experiments were approved by the Boston Children’s Hospital Institutional Animal Care and Use Committee. *Zfp410* gene-trap allele mice were generated as described below. C57BL/6 mice were obtained from Charles River Laboratories (Strain Code 027). Mouse ES cells heterozygous for a *Zfp410* gene-trap allele produced in the EUCOMM (European Conditional Mouse Mutagenesis Program) were purchased from the European Mouse Mutant Cell Repository (EuMMCR, Germany). The ES cells were derived from a C57BL/6N background. The targeting cassette was inserted in intron 5 and contains a splice acceptor site upstream of the *lacZ* gene that disrupts normal splicing and thus expression of *Zfp410* (**Supplementary Fig. 4b**). This allele also has conditional potential with LoxP sites flanking exon 6. We purchased 3 ES cell clones (E06, F06 and F07). Karyotype analysis was performed by EuMMCR. The percentage of cells with normal chromosome count (2n=40) for each clone was 77% for E06, 70% for F06 and 90% for F07. Clones E06 and F07 were chosen for blastocyst micro-injections. Chimeric mice were generated by the NIH/NIDDK Center of Excellence in Molecular Hematology, Mouse Embryonic Stem Cell (ESC) and Gene Targeting Core facility. C57BL/6 mice were used as the host for blastocyst micro-injections. For clone E06, there were a total of 13 pups born from 3 pregnant fosters, of which there were 2 male and 1 female chimeric mice. For clone F07, there were 3 pups born from 2 pregnant fosters, none of which were chimeras. Of the 3 chimeric mice obtained, one male produced germline transmission of the *Zfp410* gene-trap (Gt) allele upon breeding with wildtype C57BL/6 mice. Mice heterozygous for the *Zfp410* gene-trap allele (*Zfp410* +/Gt) were intercrossed to generate mice homozygous for the *Zf410* gene-trap allele (*Zfp410* Gt/Gt). Mice carrying the *Zfp410* gene-trap allele were genotyped using primers flanking the LoxP site in intron 6 (LoxP-FP and LoxP-RP). During homologous recombination, the targeting cassette replaces endogenous DNA stretches resulting in a slightly different PCR product from the targeted compared to the wildtype allele. Peripheral blood was collected from mice at 3 months of age. CBCs were performed on the Advia hematology system at the BCH-HSCI Flow Core. Values for the normal range of various hematological parameters for C57BL/6 mice were obtained from the Charles River Laboratories website (https://animalab.eu/sites/all/pliki/produkty-dopobrania/Biochemistry_and_Hematology_for_C57BL6NCrl_Mouse_Colonies_in_North_American_for_January_2008_December_2012.pdf). RNA was isolated from whole blood using Trizol following the manufacturer’s protocol.

### Xenotransplant

NOD.Cg-KitW-41J Tyr + Prkdcscid Il2rgtm1Wjl (NBSGW) mice were obtained from Jackson Laboratory (Stock 026622). CD34+ HSPCs from adult mobilized peripheral blood from de-identified healthy donors were thawed and recovered in X-VIVO complete medium for 24 hours. After recovery, cells were electroporated using the Lonza 4D nucleofector with 3xNLS-Cas9 protein and sgRNA. Cells were allowed to recover from electroporation for 24-48 hours in X-VIVO complete medium. Cells were counted and divided equally among 3 or 4 recipient mice per condition. A portion of cells was subjected to *in vitro* erythroid differentiation. Pre-transplant editing efficiency was assessed on day 4 of *in vitro* culture. In each experiment 4 mice received cells that were not subjected to electroporation (mock) as experimental controls. Cells were resuspended in 200 ul DPBS per mouse and infused by retro-orbital injection into non-irradiated NBSGW female mice. 16 weeks post transplantation, mice were euthanized, bone marrow was collected and xenograft analysis was performed as previously described^21^. Analysis of bone marrow subpopulations was performed by flow cytometry. Bone marrow cells were first incubated for 15 minutes with Human TruStain FcX (BioLegend #422302) and TruStain FcX (anti-mouse CD16/32, BioLegend #101320) to block non-specific binding of immunoglobulin to Fc receptors, followed by incubation with anti-human CD45 (V450, clone HI30, BD Biosciences #560367), anti-mouse CD45 (PE-eFluor 610, clone 30-F11, Thermo Fisher #61-0451-82), anti-human CD235a (FITC, BioLegend #349104), anti-human CD33 (PE, BioLegend #366608), anti-human CD19 (APC, BioLegend #302212), anti-human CD3 (PE/Cy7, BioLegend #300420) and anti-human CD34 (FITC, BioLegend #343504) antibodies. Fixable Viability Dye (eFluor 780, Thermo Fisher #65-0865-14) was used to exclude dead cells. The percentage of human engraftment was calculated as hCD45+ cells/(hCD45+ cells + mCD45+ cells) x 100. B-lymphocyte (CD19+), granulocyte (CD33-dim SSC-high) and monocyte (CD33-bright SSC-low) lineages were gated on the hCD45+ population. HSPCs (CD34+) and T-lymphocyte (CD3+) lineages were gated on the hCD45+ hCD19-hCD33-population. Human erythroid cells (CD235a+) were gated on the hCD45-mCD45-population. The detailed gating strategy is shown in **Supplementary Fig. 6**.

### Statistical analyses

All values indicated for replicates (n=x) are biological replicates. p-values were calculated by two-tailed Student’s t-test.

## Data availability

The datasets generated during the current study are available from the indicated repositories where applicable or are included in this article.

## Code availability

The scripts used for analysis of CUT&RUN experiments and motif mapping have been provided in Supplementary Methods.

## Supplementary Methods

### CUT&RUN

Sequencing data obtained from CUT&RUN experiments were analyzed using the following scripts. The workflow was largely adapted from previous protocols^9,13^, with the addition of data filtering based on findings of the ENCODE project^53,57^.

The prerequisite Software used in our methods are listed below:

FastQC 0.11.3
Trimmomatic 0.36
Bowtie 2 2.2.9
Samtools 1.3.1
Picard 2.8.0
Bedtools 2.27.1
Deeptools 3.0.2
Macs2 2.1.1.20160309

1. FastQC (Babraham Institute, version 0.11.3) was performed for all samples to check sequencing quality. $zcat *.fastq.gz $fastqc Sample1_Read1.fastq $fastqc Sample1_Read2.fastq
2. Adapter sequences were trimmed with Trimmomatic^49^. $java -jar trimmomatic-0.36.jar PE -threads 4 -trimlog trim.log -phred33 Sample1_Read1.fastq.gz Sample1_Read2.fastq.gz -baseout path-for-output/Sample1_trimmed.fq.gz ILLUMINACLIP:/adapters/TruSeq3-PE.fa:2:15:4:4:true SLIDINGWINDOW:4:15 MINLEN:25
3. Trimmed reads were aligned to either the human reference genome hg19 or the mouse reference genome mm10 using bowtie2^50^. Pre-built bowtie2 index files are available at http://bowtie-bio.sourceforge.net/tutorial.shtml. $BOWTIE2_IDX=bowtie2_indexes/hg19 (or bowtie2_indexes/mm10) $bowtie2 --end-to-end --no-unal --no-mixed --no-discordant --dovetail --phred33 -p 4 -x ${BOWTIE2_IDX} −1 Sample1_trimmed_1P.fq.gz −2 Sample1_trimmed_2P.fq.gz -S Sample1.sam
4. The resulting alignment files (.sam) were converted to sorted, indexed bam files and marked for duplicates using Picard (https://broadinstitute.github.io/picard/). $java -jar picard-2.8.0.jar SortSam I=Sample1.sam O= Sample1.sorted.bam SORT_ORDER=coordinate CREATE_INDEX=true $java -jar picard-2.8.0.jar MarkDuplicates I=Sample1.sorted.bam O=Sample1.dedup.sorted.bam M=Sample1.dedup.txt REMOVE_DUPLICATES=true $bedtools bamtobed -i $sampleID.dedup.sorted.bam > $sampleID.dedup.sorted.bed $cat Sample1.dedup.sorted.bed | awk -v OFS=‘\t’ ‘{len = $3 - $2; print $0, len }’ > Sample1.dedup.sorted.final.bed
5. Deduplicated and sorted bam files were converted to bigwig files for visualization in IGV. These are the files used to generate the representative IgG or ZNF410 data tracks. $samtools index Sample1.dedup.sorted.bam $bamCoverage --bam Sample1.dedup.sorted.bam -o Sample1.dedup.sorted.bw --binSize 10 Reads were filtered using an alignment score cutoff of 10 with samtools^51^. $samtools view -b -q 10 Sample_1.dedup.sorted.bam > Sample_1.dedup.filtered.bam
6. Peak calling was performed using macs2^52^. Biological replicates for test and control samples were grouped together at this stage. $sampleID=Sample $controlID=Control $outputI D=Sample_vs_Control $macs2 callpeak -f BAMPE -t ${sampleID}_Replicate-1.dedup.filtered.bam ${sampleID}_Replicate-2.dedup.filtered.bam ${sampleID}_Replicate-3.dedup.filtered.bam \ -c ${controlID}_Replicate-1.dedup.filtered.bam ${controlID}_Replicate-2.dedup.filtered.bam ${controlID}_Replicate-3.dedup.filtered.bam \ -B -g hs -q 0.05 -n ${outputID}
7. Genomic regions annotated as part of the ENCODE project blacklist^53,57^ as problematic regions for alignment of high-throughput functional genomics data were excluded from analysis using files ENCFF001TDO (hg19, Birney lab, EBI) and ENCFF547MET (mm10, Kundaje lab, Stanford) and BEDtools^54^. $bedtools intersect -a Test_vs_control_peaks.bed -b blacklist.bed -v > Test_vs_control_blacklist-filtered.bed

## Acknowledgements

We thank Drs. Ryo Kurita and Yukio Nakamura for sharing HUDEP-2 cells (Department of Research and Development, Central Blood Institute, Blood Service Headquarters, Japanese Red Cross Society, Tokyo, Japan and Cell Engineering Division, RIKEN BioResource Research Center, Faculty of Medicine, University of Tsukuba, Ibaraki, Japan); Dr. Ronald Mathieu and the HSCI-BCH Flow Cytometry Facility, supported by the Harvard Stem Cell Institute and the NIH (U54DK110805) for assistance with flow cytometry; Dr. Zachary Herbert from the Molecular Biology Core Facilities at Dana-Farber Cancer Institute for assistance with sequencing; Dr. Yuko Fujiwara from the BCH Mouse Embryonic Stem Cell and Gene Targeting Core (supported by the NIH/NIDDK Center of Excellence in Molecular Hematology U54DK110805) for assistance with transgenic mouse generation; Dr. John Doench for assistance with CRISPR screening; Dr. Steven Henikoff for sharing pA-MNase for CUT&RUN experiments; Dr. Scot Wolfe for sharing 3xNLS-SpCas9; Jasmine Bonanno for technical assistance; and Drs. Stuart Orkin, Christian Brendel, Nan Liu, Davide Seruggia, Neekesh Dharia, and members of the Bauer laboratory for helpful discussions. D.S.V. was supported by the Cooley’s Anemia Foundation Research Fellowship award (2018-2020); L.P. was supported by NHGRI (R00HG008399 and R35HG010717); D.E.B was supported in part by a Sponsored Research Agreement from Sanofi, NHLBI (DP2HL137300 and P01HL032262), and the Burroughs Wellcome Fund.

## Supplementary Figure Legends

**Supplementary Figure 1.**
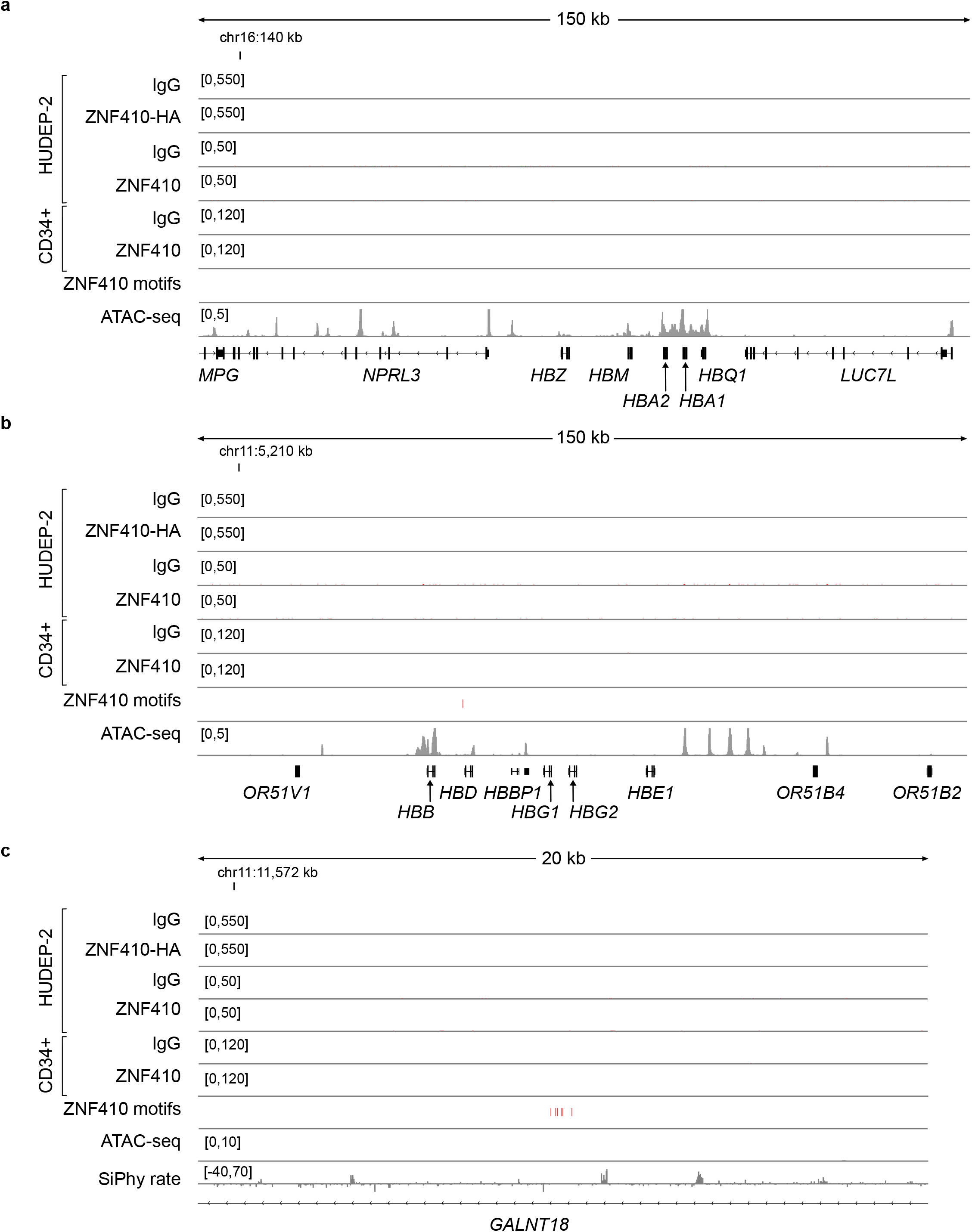
Absent ZNF410 chromatin occupancy. (a-c) α-like and *β*-like globin gene clusters and *GALNT18* intron 1 with a cluster of 6 ZNF410 motifs indicating absence of ZNF410 occupancy in representative CUT&RUN control IgG (n=9) and anti-HA (n=7) in HUDEP-2 cells over-expressing HA-tagged ZNF410, control IgG (n=1) and anti-ZNF410 (n=1) in HUDEP-2 cells, and control IgG (n=2) and anti-ZNF410 (n=2) in CD34+ HSPC derived erythroid precursors. Positions of ZNF410 motifs (red rectangles), accessible chromatin by representative ATAC-seq in HUDEP-2 cells (gray peaks, n=3) and DNA sequence conservation by SiPhy rate.

**Supplementary Figure 2.**
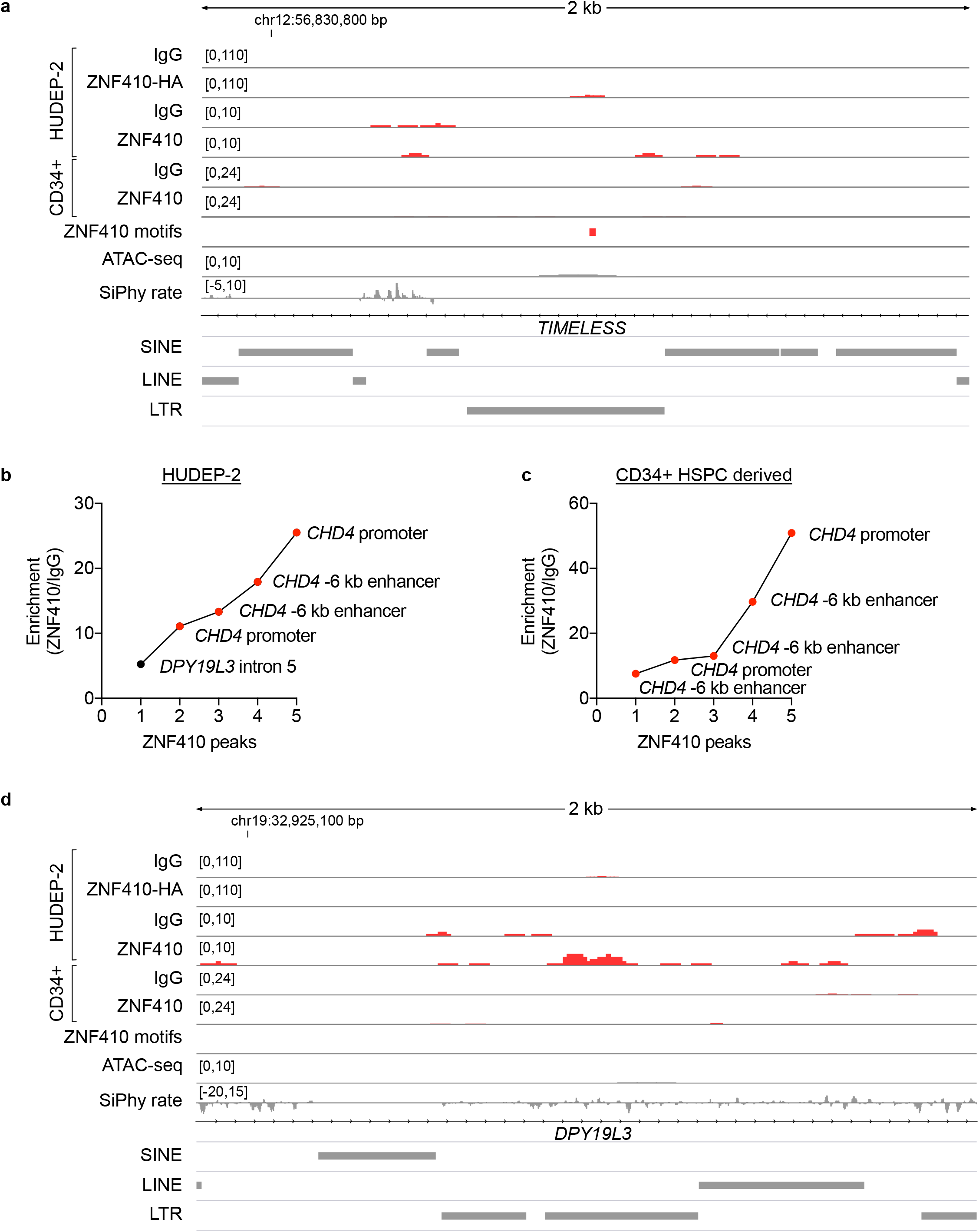
ZNF410 chromatin occupancy. (a) The third most enriched peak for ZNF410 binding (following *CHD4* promoter and −6 kb enhancer) by CUT&RUN with anti-HA antibody in HUDEP-2 cells over-expressing ZNF410-HA was at *TIMELESS* intron 1. Representative CUT&RUN control IgG (n=9) and anti-HA (n=7) in HUDEP-2 cells over-expressing HA-tagged ZNF410, control IgG (n=1) and anti-ZNF410 (n=1) in HUDEP-2 cells, and control IgG (n=2) and anti-ZNF410 (n=2) in CD34+ HSPC derived erythroid precursors. Positions of ZNF410 motifs (red rectangles), accessible chromatin by representative ATAC-seq in HUDEP-2 cells (gray peaks, n=3), DNA sequence conservation by SiPhy rate, and repetitive elements from RepeatMasker. (b) A total of 5 peaks were identified by CUT&RUN with anti-ZNF410 antibody in HUDEP-2 cells. The top 4 peaks were at the *CHD4* promoter or −6 kb enhancer, the fifth was at *DPY19L3* intron 5. (c) A total of 5 peaks were identified by CUT&RUN with anti-ZNF410 antibody in CD34+ HSPC derived erythroid precursors. All 5 peaks were at the *CHD4* promoter or −6 kb enhancer. (d) Peak of ZNF410 occupancy at *DPY19L3* intron 5 in HUDEP-2 cells.

**Supplementary Figure 3.**
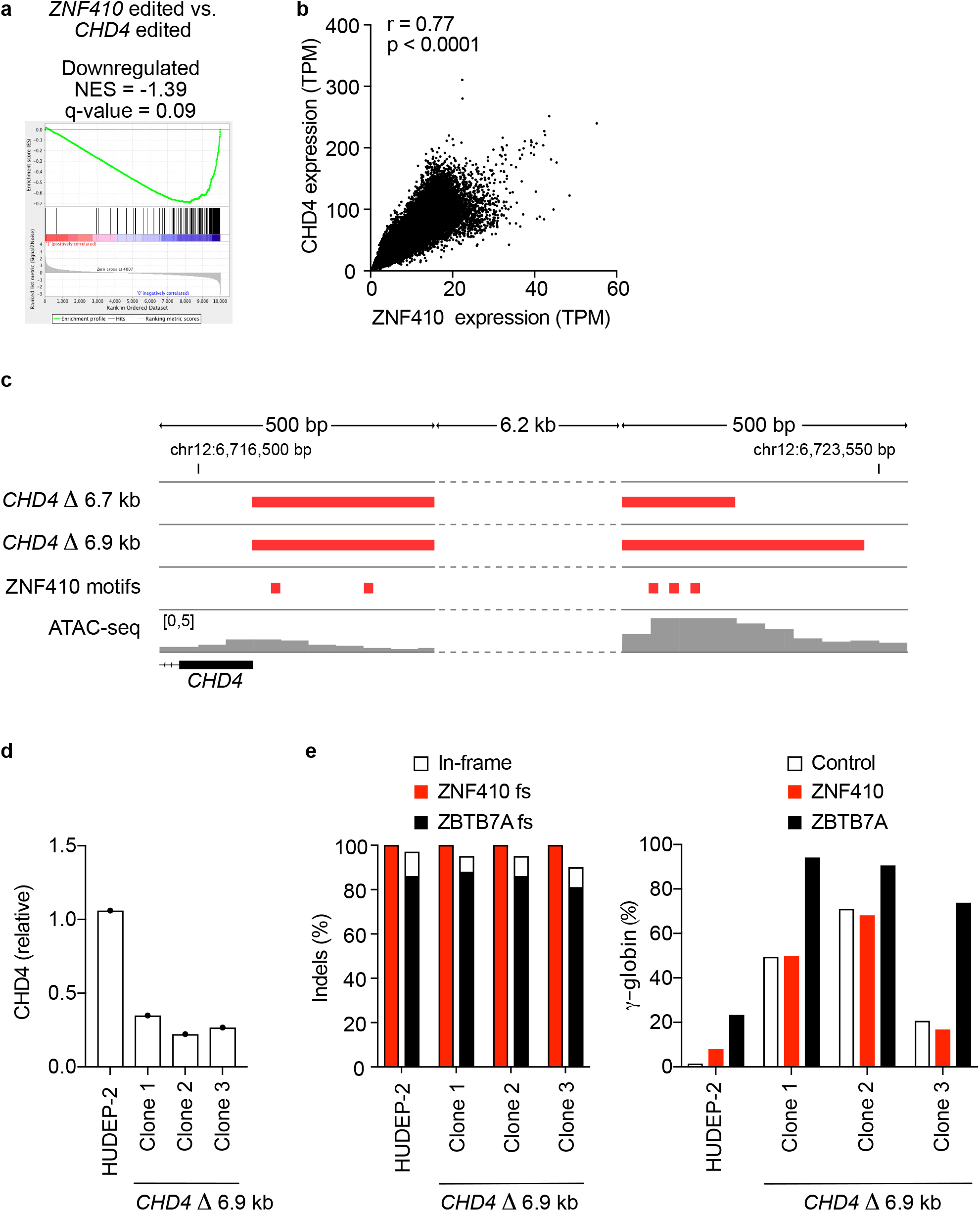
ZNF410 represses HbF by activating *CHD4*. (a) Comparison of genes downregulated in *ZNF410* and *CHD4* mutant cells by GSEA shows enrichment of *CHD4* regulated genes in the *ZNF410* regulated gene set. (b) Correlation of *ZNF410* and *CHD4* expression across 54 human tissues from GTEx (Pearson r=0.77, p<0.0001). (c) Cas9 paired cleavages with CHD4-proximal-gRNA-1 and CHD4-distal-gRNA-1 (*CHD4* Δ 6.7 kb) or CHD4-proximal-gRNA-1 and CHD4-distal-gRNA-2 (*CHD4* Δ 6.9 kb) were used to generate HUDEP-2 clones with biallelic deletions spanning both of the ZNF410 binding regions upstream of *CHD4*. Positions of ZNF410 motifs (red rectangles) and accessible chromatin by ATAC-seq (gray peaks) shown. (d) CHD4 expression measured by RT-qPCR in *CHD4* Δ 6.9 kb clones compared to HUDEP-2 cells. (e) *CHD4* Δ 6.9 kb clones and HUDEP-2 cells were subjected to *AAVS1* (negative control), *ZNF410* and *ZBTB7A* targeting using RNP electroporation of 3X-NLS-Cas9 and sgRNA. Left panel, editing efficiency measured by indel frequency in HUDEP-2 cells and *CHD4* Δ 6.9 kb clones targeted with *ZNF410* or *ZBTB7A* sgRNAs. The shaded portion of the bar represents the percentage of indels resulting in frameshift (fs) alleles. The white portion of the bar represents in-frame indels. Right panel, *HBG* expression relative to total *β*-like globin (*HBG*+*HBB*) was measured by RT-qPCR in HUDEP-2 cells and *CHD4* Δ 6.9 kb clones targeted with *AAVS1* (negative control), *ZNF410* or *ZBTB7A* sgRNAs.

**Supplementary Figure 4.**
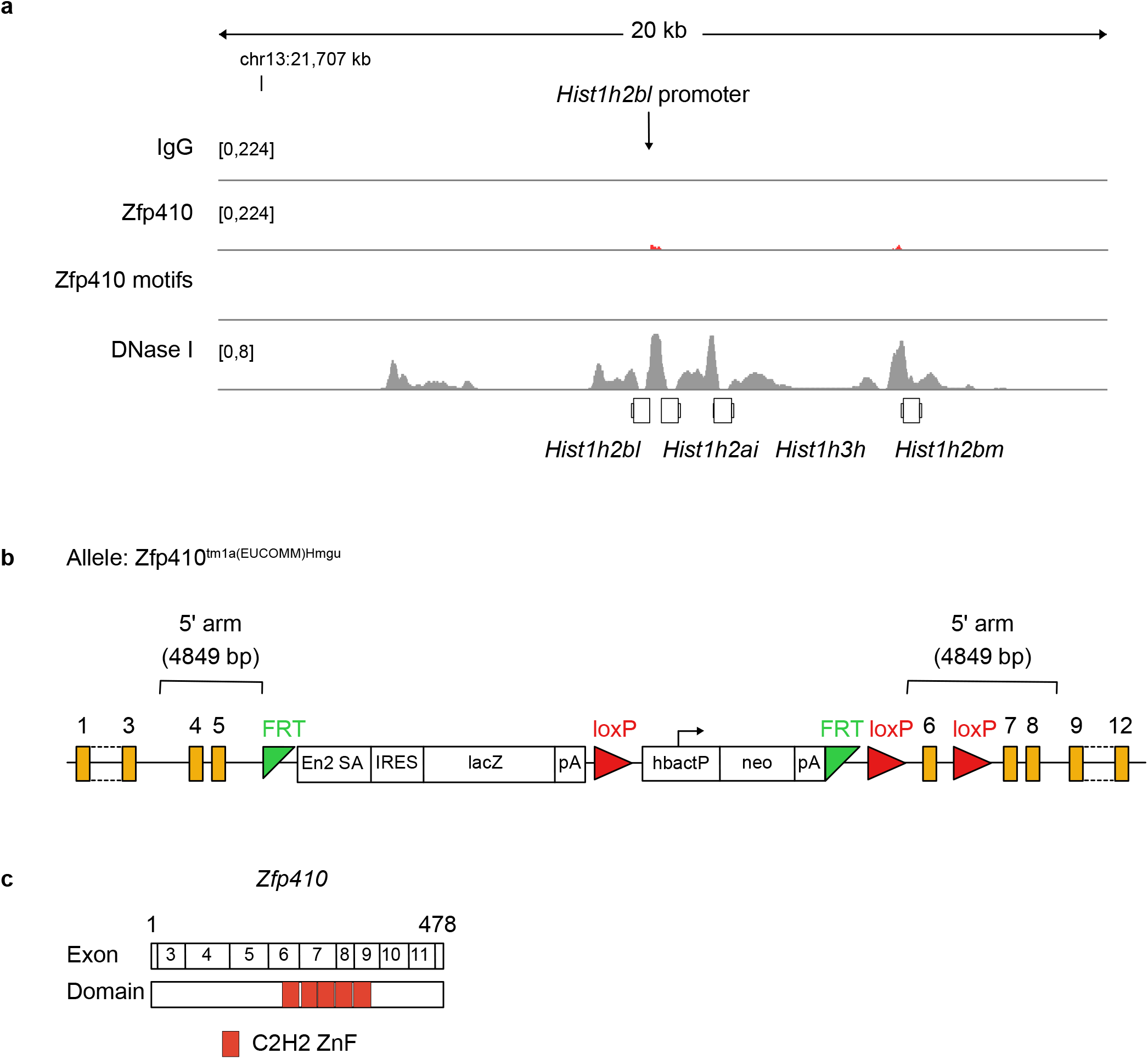
*Zfp410* is the conserved mouse ortholog of *ZNF410*. (a) CUT&RUN performed in mouse erythroleukemia (MEL) cells using anti-Zfp410 antibody (n=3) and IgG control (n=3). The third most enriched Zfp410 peak (following *Chd4* promoter and *Chd4* −6 kb enhancer) was at the *Hist1h2bl* promoter. No Zfp410 motifs were identified at this locus, which overlaps accessible chromatin (DNase-seq, gray peaks). (b) Diagram of the *Zfp410* gene trap allele. A targeting cassette including splice acceptor site upstream of *LacZ* was inserted into *Zfp410* intron 5 thus disrupting full-length expression. Schema obtained along with mouse ES cells from EuMMCR, Germany. (c) Exon and domain structure of mouse *Zfp410*.

**Supplementary Figure 5.**
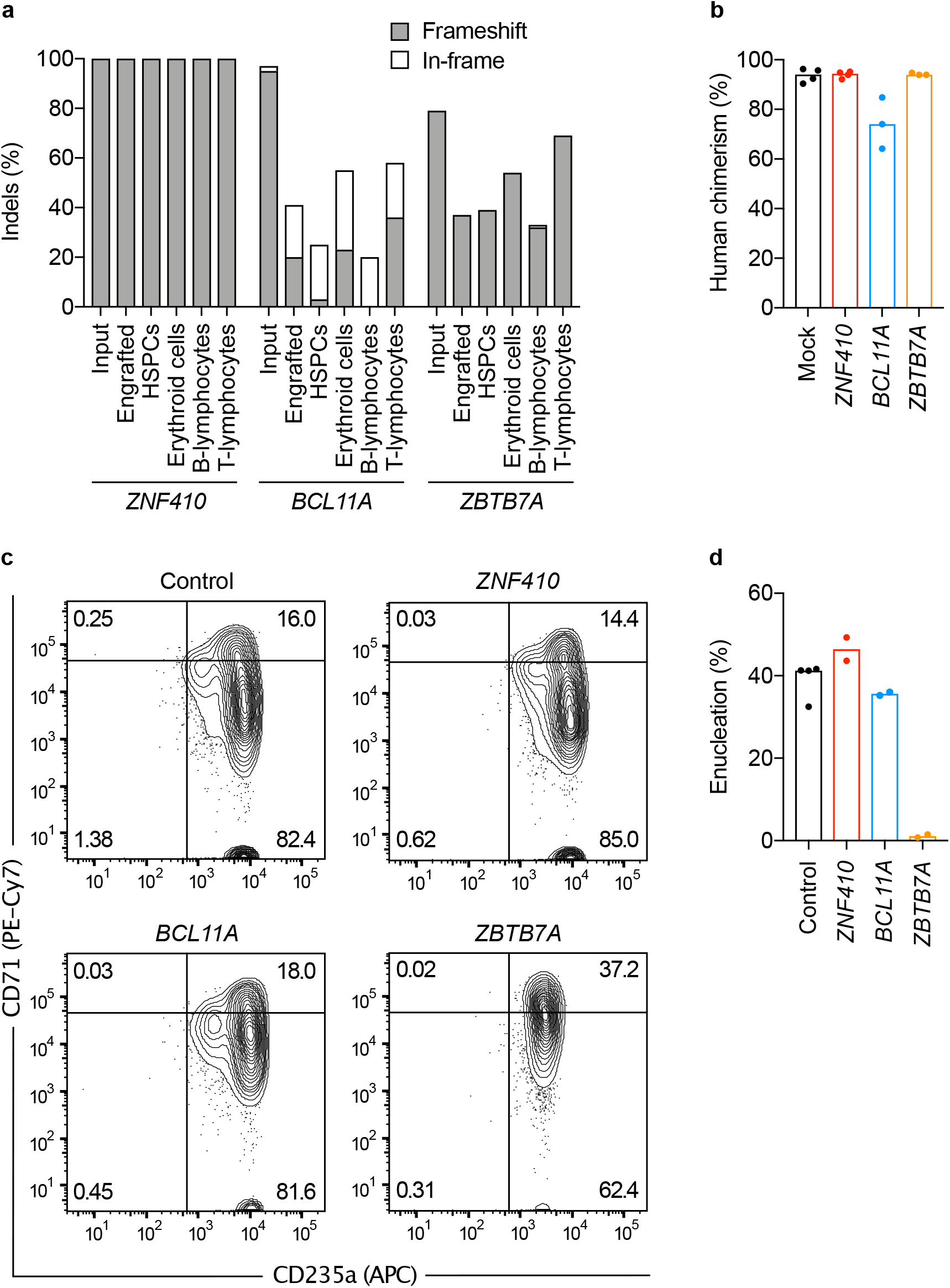
ZNF410 is dispensable for hematopoietic repopulation and erythropoiesis. CD34+ HSPCs from donor 3 were edited by RNP electroporation targeting *ZNF410, BCL11A* or *ZBTB7A* and infused to NBSGW mice or subject to in vitro erythroid differentiation. (a) Indel frequency at *ZNF410, BCL11A* and *ZBTB7A* was quantified in input cells 4 days after electroporation, and in engrafted total or sorted cells at bone marrow (BM) harvest. The percentage of frameshift alleles is represented in gray and the percentage of in-frame alleles is represented in white. (b) Comparison of engraftment assessed by human CD45+ staining compared to total CD45+ cells in xenografts of *ZNF410* (n=4), *BCL11A* (n=3) and *ZBTB7A* (n=3) edited and mock control (n=4) CD34+ HSPCs. Each symbol represents one mouse. (c, d) Erythroid maturation, evaluated based on CD71 and CD235a immunophenotype and enucleation frequency, was assessed on day 18 of *in vitro* erythroid culture.

**Supplementary Figure 6.**
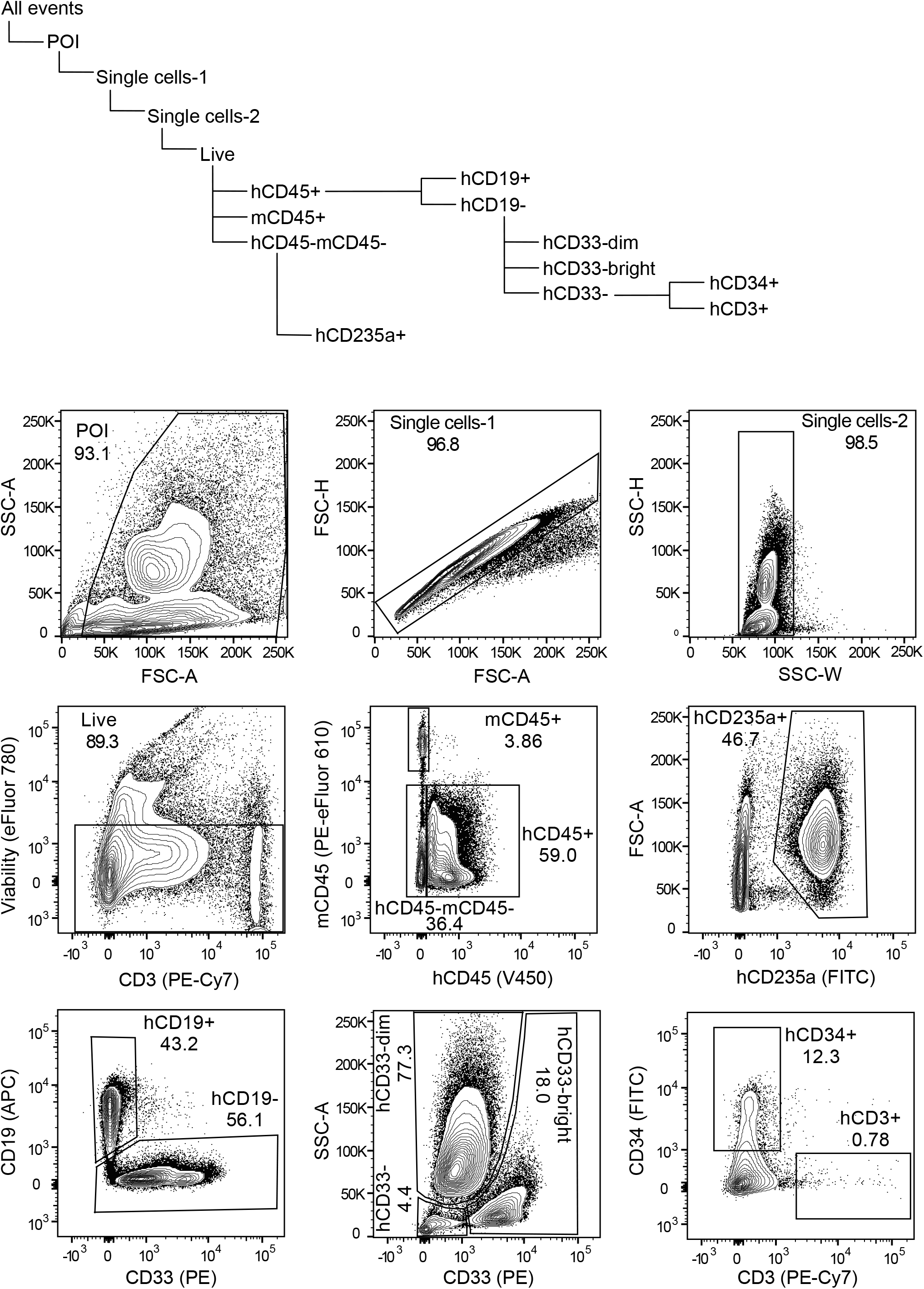
Flow cytometry gating strategy for xenograft experiment. Hierarchy of FACS gates and representative plots for each gate are shown for a representative control (mock) transplanted bone marrow sample. The first gate was plotted to delineate the cell population of interest (POI) and avoid debris. The second and third gates were plotted to exclude doublets. Values in plots are for respective gates.

---

**Supplementary Table 1.**
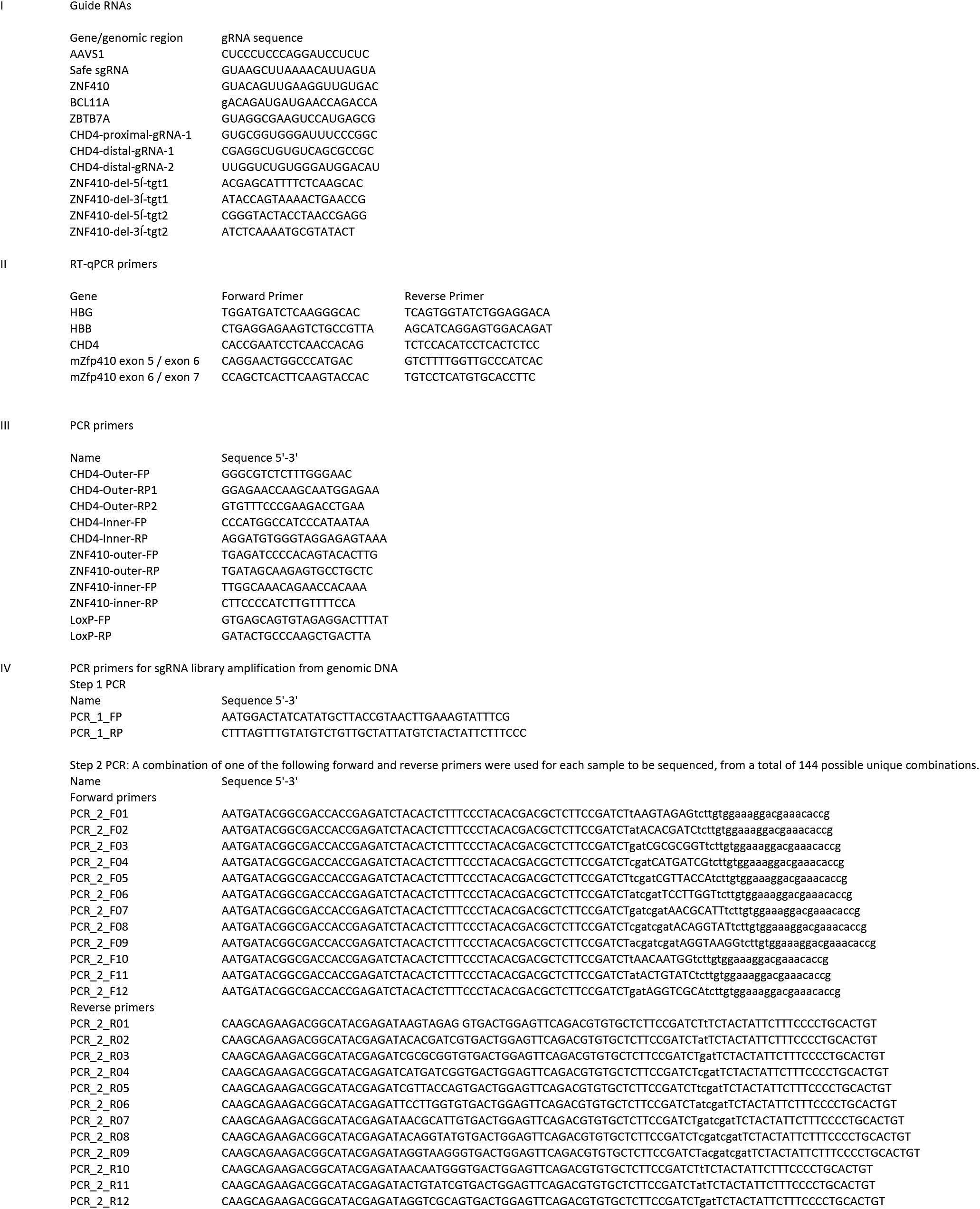

